# Dual agonism of sodium iodide symporter function *in vivo*

**DOI:** 10.1101/2024.02.27.582332

**Authors:** Katie Brookes, Caitlin M. Thornton, Ling Zha, Jana Kim, Benjamin Small, Selvambigai Manivannan, Hannah R. Nieto, Holly Adcock, Giovanni Bottegoni, Liam R. Cox, Vinodh Kannappan, Weiguang Wang, Caroline M. Gorvin, Sissy Jhiang, Matthew D. Ringel, Moray J. Campbell, Kavitha Sunassee, Philip J. Blower, Kristien Boelaert, Vicki E. Smith, Martin L. Read, Christopher J. McCabe

**Author notes:** **Corresponding Authors**: Professor Christopher J. McCabe, Institute of Metabolism and Systems Science, Birmingham Health Partners, College of Medicine and Health, University of Birmingham, Birmingham, B15 2TH, UK.; and Martin L. Read.

## Abstract

New approaches are urgently needed to enhance the radioiodide (RAI) ablation of aggressive and metastatic thyroid cancer. We recently discovered that valosin-containing protein inhibitors (VCPi) such as clotrimazole and disulfiram transiently block sodium iodide symporter (NIS) proteasomal degradation, hence promoting RAI uptake. However, poor bioavailability diminishes their potential impact *in vivo*. Following 3D modelling and iterative drug design we appraised 26 novel analogues of clotrimazole, as well as albumin nano-encapsulated copper-diethyldithiocarbamate [Cu(DDC)_2_-alb] – a stabilised reformulation of a disulfiram metabolite. While several clotrimazole analogues specifically increased RAI uptake, the greatest impact was observed with Cu(DDC)_2_-alb in thyroid cancer cells as well as human primary thyrocytes from patients with thyroid hyperplasia. NanoBRET assays revealed that Cu(DDC)_2_ enhanced the plasma membrane accumulation of NIS in living cells. In BALB/c mice, both intraperitoneal and intravenous administration of Cu(DDC)_2_-alb significantly enhanced thyroidal ^99m^Tc-uptake. RNA-Seq revealed the surprising observation that Cu(DDC)_2_-alb induced key thyroid transcription factors. Accordingly, expression of PAX8 and NKX2.1 was upregulated in thyroid glands from drug treated mice, with NIS levels correlating closely to ^99m^Tc-uptake. As Cu(DDC)_2_ inhibits the VCP cofactor NPL4, with VCP being critical to the proteostatic processing of NIS protein, we identify a new dual agonist of RAI uptake *in vivo*, with the potential to directly impact RAI therapy for patients with aggressive thyroid cancer.

## Introduction

Radioiodide treatment remains the frontline therapy for thyroid cancer, but is hampered by diminished clinical utility in subsets of patients, particularly those with aggressive and metastatic disease [1]. Recent estimates for 2022 indicated that > 47,500 lives are lost worldwide to thyroid cancer per annum, which is estimated to rise to 85,532 by 2045 [2]. New approaches and insights are hence urgently required to enhance radioiodide uptake and tumour/ metastatic ablation. Combinatorial targeting of the MAPK pathways which generally drive tumourigenesis (BRAF and MEK inhibitors) has shown recent promise in overcoming radioiodine refractoriness [3, 4], although issues of drug resistance and adverse events are common [5, 6]. Thus, the exploitation of new mechanistic insight into the function of the sodium iodide symporter (NIS), the only known conduit of iodide into mammalian cells, has the potential to transform patient therapy.

In identifying new drug approaches to increase NIS function in thyroid cells, we recently identified a panel of novel FDA-approved compounds which significantly enhanced radioiodide uptake across a variety of cell models, including human and mouse primary thyrocytes *ex vivo* [7]. A leading drug candidate was disulfiram, a proteasomal inhibitor and compound of the dicarbamate family [8], which has been approved by the US FDA since 1951 for the treatment of chronic alcoholism [9]. Mechanistic insight provided by mass spectrometry to identify novel interacting proteins of NIS also previously led us to identify the key Endoplasmic Reticulum Associated Degradation (ERAD) protein VCP/p97 as being critical to NIS protein processing [10]. A critical cofactor of VCP is the protein NPL4, and disulfiram was recently shown to inhibit NPL4 activity via its copper bound diethyldithiocarbamate metabolite Cu(DDC)_2_ [11]. We hence hypothesised that disulfiram increases radioiodide uptake by transiently interfering with ERAD via a VCP/NPL4 pathway, permitting more NIS protein to be trafficked to the plasma membrane.

A second FDA-approved drug identified in our screening approaches was the anti-fungal agent clotrimazole which potently enhanced radioiodide uptake *in vitro* [7, 10] and has been identified as a re-purposed small molecule capable of allosterically inhibiting VCP activity [12]. However, although well tolerated as an FDA-approved topical anti-fungal, clotrimazole is poorly bioavailable [12].

In the current investigation we conjectured that disulfiram and clotrimazole might represent key starting points for developing strategies to enhance radioiodide uptake *in vivo*: disulfiram and/or its metabolites might inhibit activity of the key VCP cofactor NPL4, while clotrimazole may inhibit VCP directly, with the potential to transiently block the ERAD-processing of NIS, hence increasing plasma membrane NIS localisation and uptake function. Thus, we aimed to gain a mechanistic understanding of how disulfiram and its DDC metal complex Cu(DDC)_2_ impact NIS function in thyroid cells *in vitro* and *in vivo*, and whether we might be able to overcome the poor bioavailability of clotrimazole. We show that while computationally designed clotrimazole analogues enhanced radioiodide uptake *in vitro*, the disulfiram metabolite Cu(DDC)_2_ was more tractable *in vivo*. Contrary to its cardinal function of inhibiting NPL4, Cu(DDC)_2_ did not modulate NIS function via this route. Instead, RNA-Seq analysis revealed that Cu(DDC)_2_ induced the expression of key NIS transcription factors, including PAX8. In BALB/c mice, intraperitoneal (IP) and intravenous (IV) administration of physiological doses of Cu(DDC)_2_ were accompanied by significant increases in ^99m^Tc-uptake in thyroid glands but not other organs, as well as significantly inducing NIS, PAX8 and NKX2-1 expression. Our findings therefore identify a promising and entirely novel pathway via which NIS function may be augmented in the clinical setting.

## Materials and Methods

### Human thyroid tissue

This study was conducted according to the Declaration of Helsinki ethical guidelines and collection of normal human thyroid tissue was approved by the Local Research Ethics Committee (Birmingham Clinical Research Office, Birmingham, UK). Informed written consent was obtained from each subject. No age/gender information was available as subjects were anonymized and tissue collected as excess to surgery as part of our ethics agreement. Primary thyrocytes were isolated and cultured as described [7].

### Animal experiments

All animal experiments were performed in accordance with the Animals (Scientific Procedures) Act, 1986 with protocols approved by the Animal Welfare and Ethical Review Body for King’s College London (St Thomas’ Campus). Male BALB/c mice (8-10 weeks of age, *n* = 5-7 animals/group, Charles River Laboratories) received either vehicle (PBS) or Cu(DDC)_2_-albumin nanoparticles (3-5 mg/kg) by either intraperitoneal (IP) or intravenous (IV) injection on days 1 and 3 of the experiment. On day 4, mice were anaesthetized by isoflurane inhalation (3%, Animalcare, York, in O_2_) and maintained under isoflurane anesthesia during IV administration of ^99m^Tc-pertechnetate (^99m^Tc; 0.5 MBq). After 30 min, mice were culled by anesthetic overdose and tissues harvested. Thyroid glands were removed using a dissecting microscope and radioactivity measured by gamma counting (1282 Compugamma; LKB Wallac). CB5083 (15-25 mg/kg/day) was given to male BALB/c mice (8-10 weeks of age, n = 3 animals/group, Charles River Laboratories) by oral gavage (OG) administration for 4 days prior to IV administration of ^99m^Tc.

### Drugs and in silico modelling

VCP inhibitors CB5083 and CB5399 were kindly provided by Cleave Therapeutics (San Francisco, USA). Cu(DDC)_2_-albumin was kindly provided by Disulfican (University of Wolverhampton). Other drugs used included Cu(DDC)_2_ (Tokyo Chemical Industry; dissolved in DMSO) and copper(II) D-gluconate (Sigma-Aldrich; dissolved in H_2_0), while compounds C22 to C25 were synthesized by SIA Enamine (Latvia) and dissolved in DMSO. Modified clotrimazole compounds (C1 to C21 and C26) were synthesized using standard procedures (School of Chemistry, University of Birmingham) and resuspended in DMSO. Predicted binding of clotrimazole to an allosteric binding site in VCP was modelled using the UCSF Chimera program [13] and 3D-coordinates from the cryo-EM resolved structure of VCP in complex with allosteric VCP inhibitor UPCDC30245 (PDB: 5FTJ) [14]. All drugs were diluted in RPMI-1640 medium (1:100; Life Technologies) prior to treatment of cells. For IP and IV administration in mice, Cu(DDC)_2_-albumin was formulated in PBS. CB5083 was resuspended in 0.5% methyl cellulose (400cp/water) using a mortar and pestle and vortexed prior to OG administration in mice.

### Cell culture

Thyroid (TPC-1, 8505C, SW1736, BCPAP) cancer cell lines were maintained in RPMI-1640 (Life Technologies), while HeLa cancer cells were maintained in DMEM (Sigma-Aldrich). Media was supplemented with 10% fetal bovine serum (FBS), penicillin (10^5^ U/l), and streptomycin (100 mg/l) and cell lines were maintained at 37°C and 5% CO_2_ in a humidified environment. Cell lines were obtained from ECACC (HeLa) and DSMZ (8505C), while TPC-1, SW1736 and BCPAP cell lines were kindly provided by Professor Rebecca Schweppe (University of Colorado). The L87-NIS cell line was kindly provided by Professor Christine Spitzweg (Ludwig-Maximilians University, Munich). Cells were cultured at low passage, authenticated by short tandem repeat analysis (NorthGene; Supp Fig. S1) and tested for mycoplasma contamination (EZ-PCR kit; Geneflow). Thawed cells were cultured for at least 2 weeks prior to use. Stable TPC-1-NIS and 8505C-NIS cells were generated by transfection of parental TPC-1 or 8505C cells with pcDNA3.1-NIS. Geneticin-resistant monoclonal colonies were expanded following FACS single cell sorting (University of Birmingham Flow Cytometry Facility), and Western blotting used to confirm NIS expression.

### Nucleic acids and transfection

To construct plasmids for NanoBiT assays VCP cDNA was cloned into pcDNA3.1 with an added N-terminal LgBiT tag, whereas a SmBiT tag was added to the C-Terminal of NIS cDNA as previously described [7]. NIS-NanoLuc (Nluc) cDNA was synthesized and subcloned into pcDNA3.1 by GeneArt (ThermoFisher Scientific) for NanoBRET detection. Professor Nevin Lambert (Georgia Regents University) kindly provided the NanoBRET plasma membrane marker Kras-Venus, as well as the Venus-tagged subcellular compartment marker RAB11. Venus-tagged markers RAB1 and RAB8 were kindly provided by Professor Kevin Pfleger (University of Western Australia) [15]. Plasmid DNA and siRNA transfections were performed with TransIT-LT1 (Mirus Bio) and Lipofectamine RNAiMAX (ThermoFisher Scientific) following standard protocols in accordance with the manufacturer’s guidelines.

### NanoBiT and NanoBRET live cell assays

Cells were seeded in 6-well plates at a density of 3.5 x 10^5^ cells per well and transfected with 500 ng – 1 µg plasmid DNA (e.g. 50 ng pcDNA3.1-NIS-Nluc and 500 ng pcDNA3.1-KRAS-Venus). 24 hr post-transfection, cells were harvested and reseeded into white 96-well plates in phenol-red-free DMEM (Life Technologies). Furimazine (Promega) was added to each well in accordance with the manufacturer’s guidelines and readings taken at 120 s intervals for up to 40 min (PHERAstar FS microplate reader; BMG Labtech). In some experiments, cells were treated with CB5339 or Cu(DDC)_2_ prior to addition of furimazine. NanoBRET signal was calculated using standard protocols by dividing the acceptor emission at 535 nm by the donor emission at 475 nm before normalizing to background signal.

### Western blotting and RAI uptake

Western blotting and RAI (^125^I) uptake assays were performed as described previously [7, 16]. Blots were probed with specific antibodies against NIS (1:1000; Proteintech), NPL4 (1:500; Cell Signaling Technology), VCP (1:1000; Cell Signaling Technology) and β-actin (1:10000; Sigma-Aldrich). HRP-conjugated secondary antibodies (Agilent Technologies) against either mouse or rabbit IgG were used at 1:2000 dilution.

### RNA-Seq and qPCR

Total RNA was extracted using the RNeasy Micro Kit (Qiagen) and reverse transcribed using the Reverse Transcription System (Promega). Mouse thyroid tissue was homogenized in buffer RLT using TissueLyser II (Qiagen; 2x 2 min cycles at 30 Hz) and 5 mm stainless steel beads. Expression of specific mRNAs was determined using 7500 Real-time PCR system (Applied Biosystems) as described previously [7]. TaqMan qPCR assays used are listed in Supplementary Information. RNA sequencing (RNA-Seq) was performed in the presence of Cu(DDC)_2_ (250 nM, 24 hr) versus vehicle in 8505C cells. RNA-seq libraries were prepared at the Next Generation Sequencing Facility (University of Birmingham). Transcript abundance estimates were normalized and differentially expressed genes (DEGs) identified using a standard edgeR pipeline, and functional classification (DAVID, ToppGene and GSEA) performed.

### Statistical analyses

Statistical analyses were performed using IBM SPSS Statistics (Version 29), GraphPad Prism (Version 9.5) and Microsoft Excel. All results were obtained from triplicate biological experiments unless otherwise indicated. For comparison between two groups, data were subjected to the Student’s t-test, and for multiple comparisons one-way ANOVA was used with either Dunnett’s or Tukey’s post-hoc test. P-values were adjusted using the Benjamini-Hochberg FDR correction procedure to correct for multiple comparisons. *P* < 0.05 was considered significant. All *P*-values reported from statistical tests were two-sided.

### Data availability

Data generated in this study are available from the corresponding author upon request.

## Results

### Developing new strategies to drug VCP inhibition of NIS function *in vivo*

Our previous drug screening identified 2 novel potential strategies to induce NIS function *in vivo*: (i) inhibition of VCP, which is critical to NIS protein processing via ERAD, and (ii) targeting general proteasomal degradation of NIS via the drug disulfiram (DSF) [7, 10]. Addressing the first approach, as outlined in Fig. 1A, the specific VCP inhibitors CB5083 and CB5339 [17, 18] induced NIS function in TPC-1 thyroid cancer and L87 cells stably expressing NIS (Fig. 1B; Supp Fig. S2A and S2B), as well as in human primary thyrocytes from multinodular goitre patients, paving the way for murine studies (Fig. 1C). However, CB5083 did not induce thyroidal uptake of ^99m^Tc in wild type BALB/c mice (Supp Fig. S2C).

**Figure 1.**
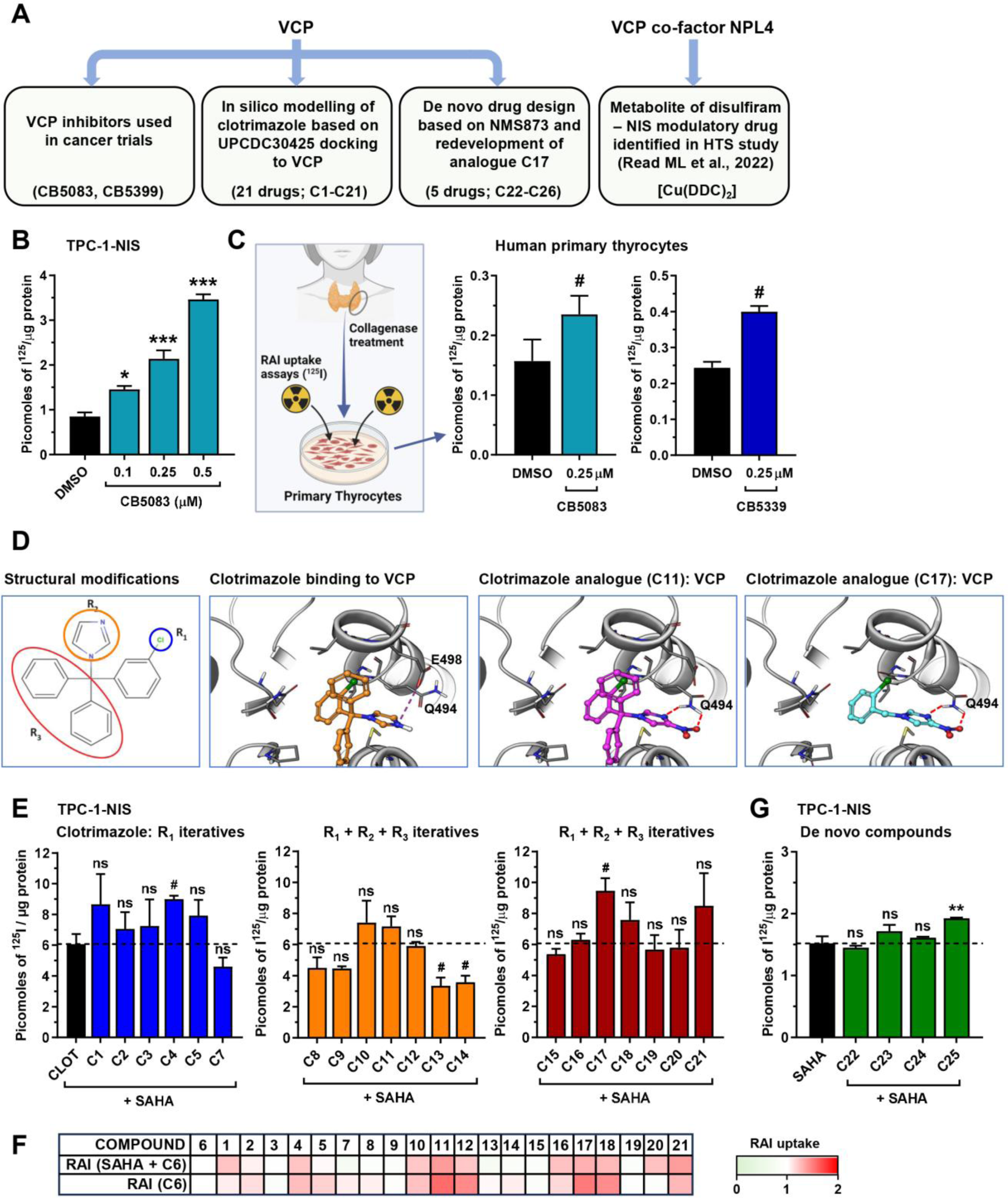
New strategies to target VCP and increase RAI uptake. **A,** Overview of drug development approaches used to target VCP or its co-factor NPL4 and enhance NIS function. **B,** RAI uptake in TPC-1-NIS cells treated with CB5083 at indicated doses for 24 hr. See also Supp Fig. S2A and S2B. **C,** RAI uptake in human primary thyrocytes treated with CB5083 or CB5339 at indicated dose. (*left*) Schematic depicting the culturing of human primary thyrocytes and RAI uptake. Created with BioRender.com. **D,** Modelling the binding of clotrimazole, compound C11 and compound C17 to VCP. Dashed lines represent predicted hydrogen bonds with residues (Q494, E498) in the allosteric binding pocket of VCP. (*left*) Chemical structure of clotrimazole highlighting modifications made at the chloro-substituted aryl ring (R_1_), imidazole ring (R_2_) and two aryl substituent groups (R_3_). **E,** RAI uptake in TPC-1-NIS cells treated with 21 compounds (C1-C21; 12 hr) in combination with SAHA (24 hr) versus clotrimazole (CLOT, C6) + SAHA. Dashed line represents normalised RAI uptake using CLOT in combination with SAHA. See also Supp Fig. S3G and Supp Table S1. **F,** Heat map depicting mean RAI uptake in TPC-1-NIS and 8505C-NIS cells treated with 21 compounds versus clotrimazole (C6) alone (*lower*) or in combination with SAHA (*upper*). See also Supp Table S1. **G,** RAI uptake in TPC-1-NIS cells treated with 4 compounds (C22-C25) in combination with SAHA versus SAHA alone. Dashed line represents RAI uptake using SAHA alone. See also Supp Fig. S4. Data presented as mean ± S.E.M., one-way ANOVA followed by Dunnett’s post hoc test (ns, not significant; \**P* < 0.05; \*\**P* < 0.01; \*\*\**P* < 0.001), or unpaired two-tailed t-test (^#^*P* < 0.05).

Whilst highly specific for VCP, CB5083 and CB5339 are competitive ATPase inhibitors; previous data [10] suggested that allosteric inhibition of VCP via the drug clotrimazole is an alternative route to drug NIS activity (Supp Fig. S2D). Clotrimazole has low bioavailability and hence we used computational iterative drug design to model and construct 21 analogues (C1-C21) with improved bioavailability, based on docking of the established allosteric VCPi UPCDC30425 to VCP (Supp Fig. S3A; Supp Table S1). Debulking of the two aryl groups (Fig. 1D; R3) in clotrimazole had the greatest overall impact on drug bioavailability and NIS activity (Supp Fig. S3B-S3F; Supp Table S1), which allowed us to construct 5 additional compounds with good predicted bioavailabilities and VCP interaction (C22-C26; Supp Fig. S4). When tested either alone or in combination with SAHA, which we have previously shown to additively increase RAI uptake with clotrimazole [7], none of the 26 compounds markedly enhanced RAI uptake to warrant taking forward for *in vivo* investigation (i.e. < 2-fold; Fig. 1E and 1F; Supp Fig. S3G and S4; Supp Table S1). We therefore confined subsequent experiments to our second strategy: targeting NIS proteasomal degradation via the proteasomal inhibitor disulfiram.

### Disulfiram metabolite Cu(DDC)_2_ potently induces RAI uptake *in vitro*

We previously identified disulfiram as a novel enhancer of RAI uptake *in vitro* [7]. However in the current study no significant increase in radionuclide accumulation was noted in the thyroid, or in any other murine organ assessed, following administration of disulfiram in wild type BALB/c mice (data not shown). Disulfiram is metabolised to diethyldithiocarbamate (DDC) *in vivo*, a hydrophilic and highly polar intermediatory which readily binds elemental copper to form the DDC-metal complex Cu(DDC)_2_ (Fig. 2A). Recently, it has been discovered that Cu(DDC)_2_ binds and inhibits the VCP co-factor NPL4 [11]. We therefore addressed the hypothesis that inhibition of VCP’s critical co-factor NPL4 – rather than VCP itself – might represent a sensitive and alternative strategy to induce NIS function *in vivo*.

**Figure 2.**
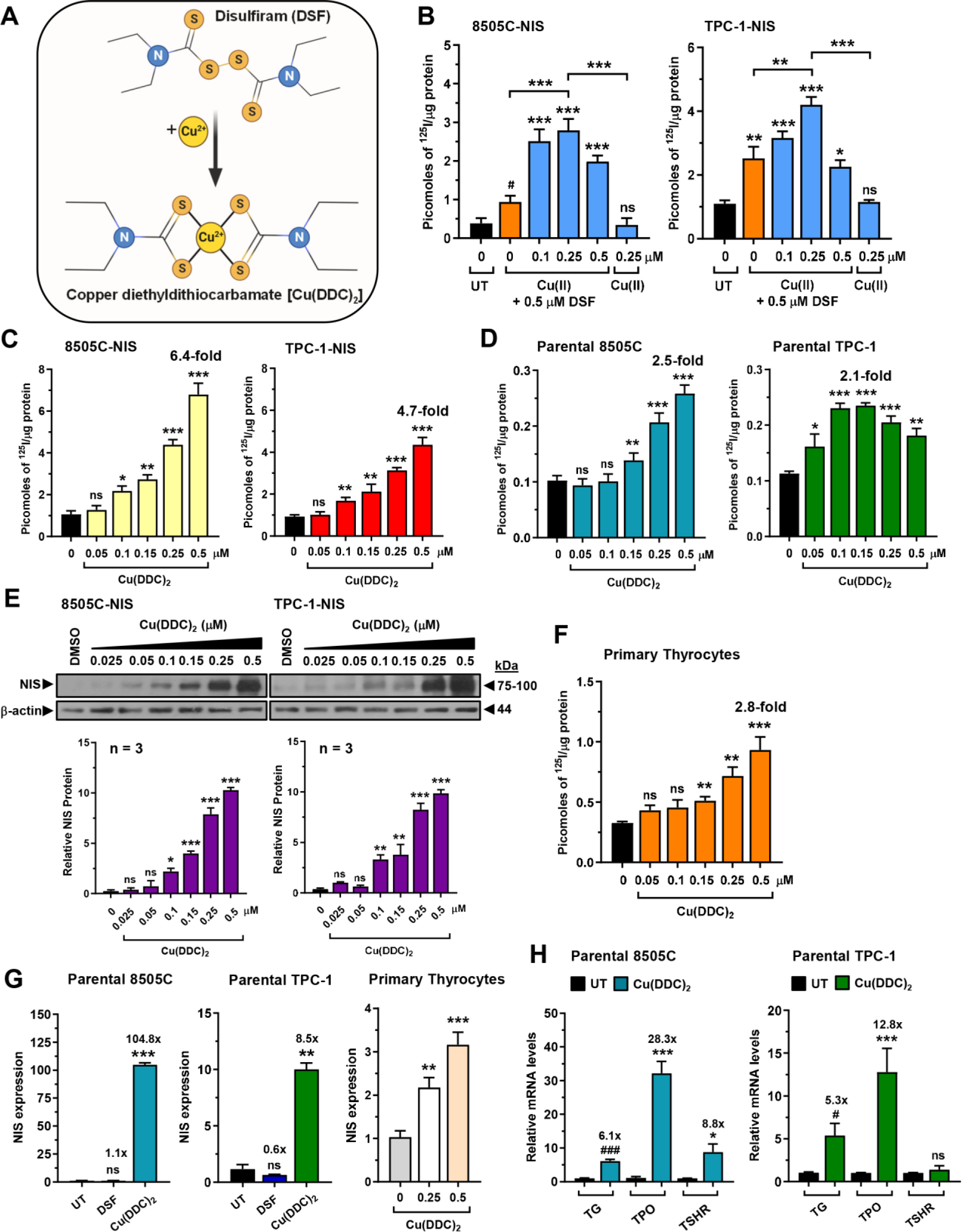
Metabolite Cu(DDC)_2_ augments NIS expression and RAI uptake in primary and cancer thyroid cells. **A,** Schematic showing metabolism of disulfiram (DSF) into copper diethyldithiocarbamate [Cu(DDC)_2_]. **B,** RAI uptake of 8505C-NIS and TPC-1-NIS cells treated with copper gluconate [Cu(II)] at indicated doses either alone or in combination with 0.5 μM DSF versus untreated (UT). **C,** RAI uptake of 8505C-NIS and TPC-1-NIS cells treated with Cu(DDC)_2_ at indicated doses for 24 hr. **D,** Same as **C** but in parental 8505C and TPC-1 cells. **E,** Western blot analysis of NIS expression in 8505C-NIS and TPC-1-NIS cells treated with Cu(DDC)_2_ at indicated doses for 24 hr. (*below*) Quantification of NIS protein levels. **F,** RAI uptake in human primary thyrocytes treated with Cu(DDC)_2_ at indicated doses. **G,** Relative NIS mRNA levels in parental 8505C and TPC-1 cells treated with DSF or Cu(DDC)_2_, as well as in human primary thyrocytes treated with Cu(DDC)_2_ at different doses. **H,** Relative TG, TPO and TSHR mRNA levels in parental 8505C and TPC-1 cells treated with Cu(DDC)_2_ for 24 hr. Data presented as mean ± S.E.M., one-way ANOVA followed by Dunnett’s or Tukey’s post hoc test (ns, not significant; \**P* < 0.05; \*\**P* < 0.01; \*\*\**P* < 0.001), or unpaired two-tailed t-test (^#^*P* < 0.05; ^###^*P* < 0.001).

*In vitro*, copper gluconate augmented disulfiram’s potentiation of RAI uptake in TPC-1 and 8505C cells stably expressing the NIS protein (Fig. 2B). Copper gluconate alone did not increase ^125^I uptake (Supp Fig. 5A). Across multiple thyroid cancer cell lines, either stably expressing NIS or not, Cu(DDC)_2_ markedly increased RAI uptake in a dose-dependent manner (Fig. 2C and 2D; Supp Fig. 5B), accompanied by parallel increases in NIS – but not VCP (Fig. 2E; Supp Fig. 5C) – protein expression. Further, Cu(DDC)_2_ was active in human primary thyrocytes (Fig. 2F), indicating that it impacts endogenous NIS activity, rather than merely overcoming neoplastic repression of NIS expression and function. Overall across all cell types, the mean maximal induction of RAI uptake via Cu(DDC)_2_ was approximately 3.4-fold.

As Cu(DDC)_2_ treatment was associated with increased NIS protein we assessed the effect of Cu(DDC)_2_ on NIS transcription, and noted a marked impact on NIS mRNA levels both in parental and stable NIS-expressing TPC-1 and 8505C cells, as well as in human primary thyrocytes (Fig. 2G; Supp Fig. S5D). In contrast, disulfiram had a much less pronounced and inconsistent impact on NIS mRNA. Unexpectedly, Cu(DDC)_2_ also increased the expression of the critical thyroid genes thyroglobulin (*TG*) and thyroid peroxidase (*TPO*; Fig. 2H). TG and TPO are central to thyroid hormone biosynthesis, frequently used as markers of thyroid cell differentiation status, and generally expressed at low levels in thyroid cancer cell lines such as 8505C and TPC-1.

### Dissecting the mechanistic impact of disulfiram metabolite Cu(DDC)_2_ in thyroid cells

To better understand the biological impact of Cu(DDC)_2_, we first investigated the cytotoxicity of Cu(DDC)_2_ in 8505C and TPC-1 thyroid cancer lines stably expressing NIS. In contrast to the methylated and inactive metabolite of disulfiram (MeDDC), which did not induce RAI uptake (Fig. 3A), Cu(DDC)_2_ exhibited a dose-dependent impact on cell viability, presumably reflecting its well described role as a proteostatic inhibitor. IC_50_ values were 38.13 and 0.37 μM in 8505C and TPC-1 cells, respectively (Fig. 3B). Our findings of cancer cell toxicity are in line with disulfiram’s reported anti-tumour properties [8].

**Figure 3.**
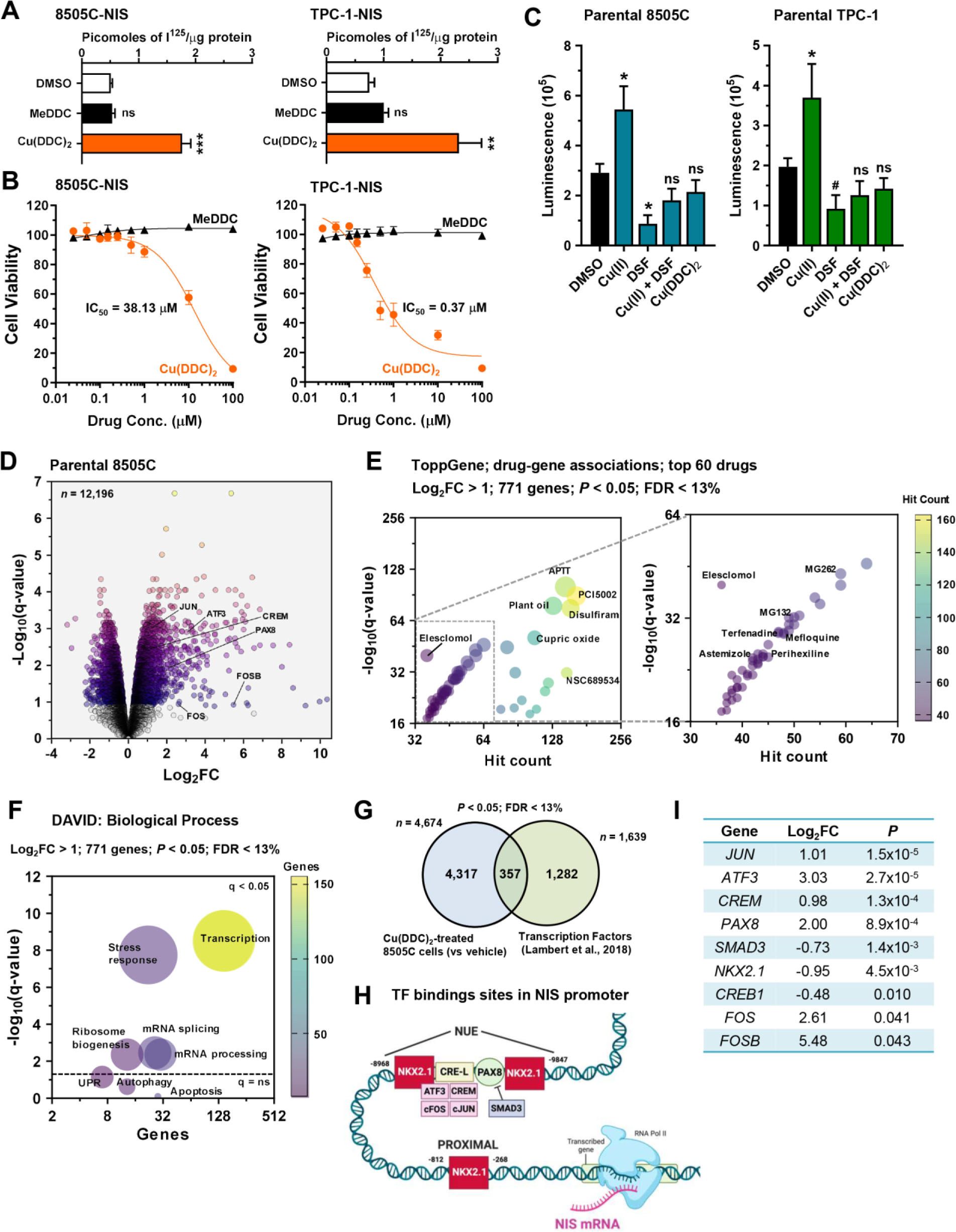
Cu(DDC)_2_ elicits a potent transcriptional response modulating NIS regulators. **A** and **B,** RAI uptake (**A**) and cell viability assays (**B**) in 8505C-NIS and TPC-1 NIS cells treated with Cu(DDC)_2_ and S-methyl N,N-diethyldithiocarbamate (MeDDC) at indicated doses for 24 hr. **C,** ROS-Glo assay evaluation of H_2_O_2_ levels in parental 8505C and TPC-1 cells treated with Cu(II), DSF, Cu(II) + DSF and Cu(DDC)_2_ versus DMSO as indicated. **D,** Volcano plot comparing log_2_FC with q-value (-log base 10) for RNA-Seq analysis (*n* = 12,196 genes) of parental 8505C cells treated with 0.25 μM Cu(DDC)_2_ versus UT. Genes associated with regulation of the NIS promoter are indicated. **E,** ToppGene classification of drug-gene associations in top 771 differentially expressed genes (log_2_FC > 1, *P* < 0.05, FDR < 13%) in parental 8505C cells treated with 0.25 μM Cu(DDC)_2_ versus UT. (*right*) enlarged view of drug-gene associations with hit count < 64. **F,** DAVID: biological process classification of top 771 differentially expressed genes (log_2_FC > 1, *P* < 0.05, FDR < 13%) as described in **D**. **G,** Venn diagram showing overlap in differentially expressed genes as described in **D** (*n* = 4,674, *P* < 0.05, FDR < 13%) versus a database of known human transcription factors [22]. **H,** Schematic showing relative position of transcription factor (TF) binding sites in the human NIS promoter. Created with BioRender.com. **I,** Candidate list of Cu(DDC)_2_-modulated NIS transcriptional regulators. Log_2_FC values are given as well as *P*-values (FDR < 13%). Data presented as mean ± S.E.M., one-way ANOVA followed by Dunnett’s post hoc test (ns, not significant; \**P* < 0.05; \*\**P* < 0.01; \*\*\**P* < 0.001), or unpaired two-tailed t-test (^#^*P* < 0.05).

In addition to inhibiting proteasomal activity, disulfiram’s anticancer capability has been reported to include elevating reactive oxygen species (ROS) levels [8, 19, 20]. We therefore tested whether disulfiram and its principal metabolites induce significant ROS in thyroid cells. Whereas copper gluconate increased ROS levels in 8505C and TPC-1 cells, disulfiram and Cu(DDC)_2_ failed to do so (Fig. 3C; Supp Fig. S5E). As copper gluconate itself did not increase cellular RAI uptake (Supp Fig. S5A), it is therefore unlikely that Cu(DDC)_2_’s mechanism of action primarily involves the induction of ROS, which is known to impact NIS function [21].

Our hypothesis was initially focussed on the alternative mechanism: that disulfiram and its Cu(DDC)_2_ metabolite inhibit the proteasomal degradation of NIS, either via VCP or NPL4. However, given the marked transcriptional changes induced by Cu(DDC)_2_ in thyroid cells (Fig. 2G and 2H), and the lack of an impact on ROS, we performed RNA-Seq analysis to identify transcriptional pathways altered by Cu(DDC)_2_ treatment (Fig. 3D). Functional classification identified strong drug: gene associations with VCP/proteasomal inhibitors (e.g. disulfiram), as well as copper-related drugs (Fig. 3E; Supp Table S2). Importantly, Cu(DDC)_2_ treatment was also associated with potent transcriptional changes in 8505C cells (Fig. 3F; Supp Fig. S5F and S5G), including significant dysregulation (*P* < 0.05; FDR < 13%) of ∼22% of all genes (357/1639) reported to encode for transcription factors in the human genome (Fig. 3G) [22].

Interestingly, we identified several transcription factors regulated by Cu(DDC)_2_ with well-characterised binding sites in the NIS promoter and key roles in regulating NIS expression (Fig. 3H and 3I) [23]. Subsequently, we confirmed via TaqMan RTPCR that multiple transcription factors, including PAX8, CREM and ATF3 (Fig. 4A; Supp Fig. 6), were all significantly induced by drug treatment compared with controls. Thus, apart from its described role as an inhibitor of the VCP co-factor NPL4 [11], we have identified a new role for Cu(DDC)_2_ as a transcriptional regulator, with a potential dual agonist role in RAI uptake.

**Figure 4.**
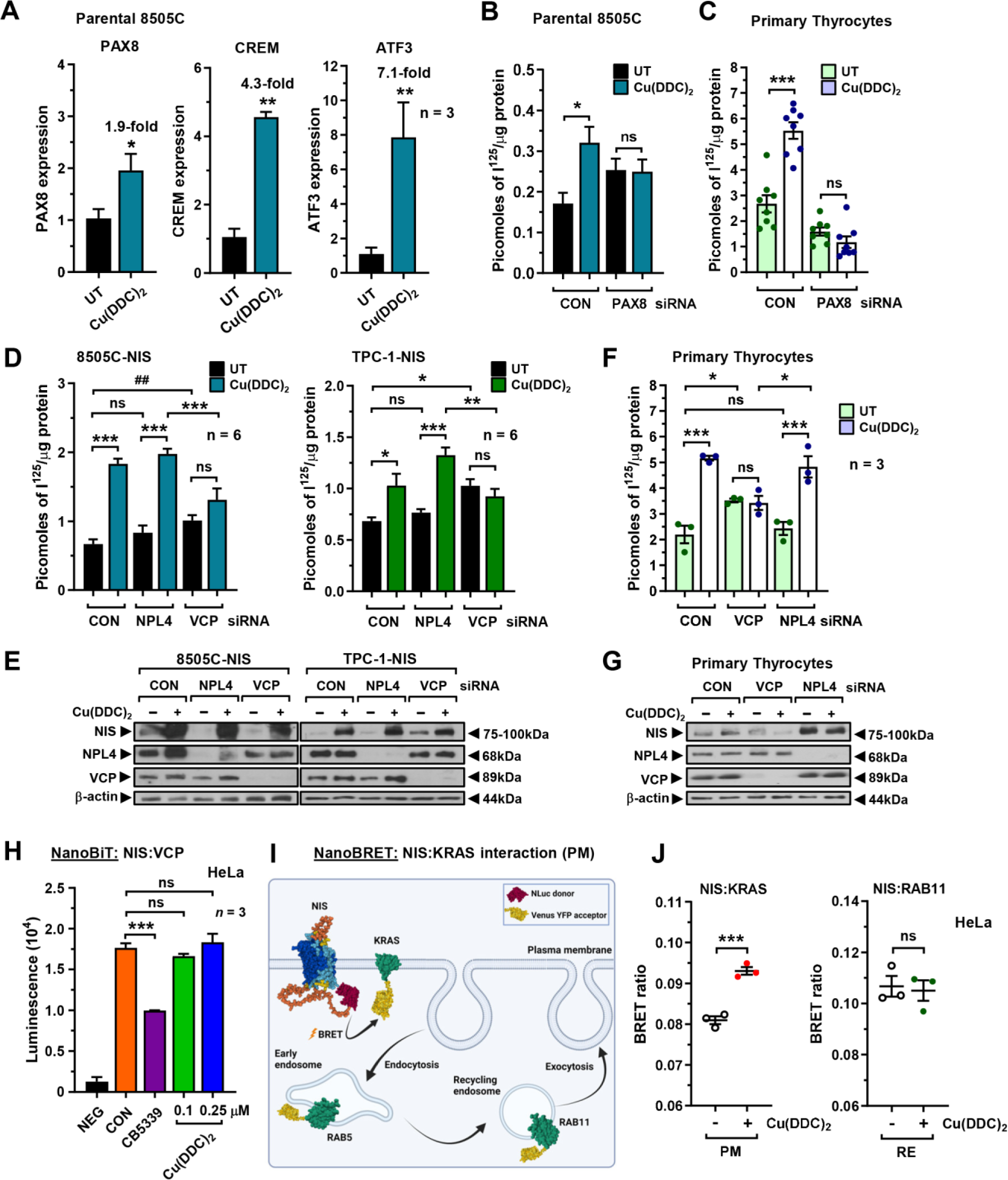
Enhanced NIS function by Cu(DDC)_2_ is dependent on PAX8 and VCP but not NPL4. **A,** Relative PAX8, CREM and ATF3 mRNA levels in parental 8505C cells treated with Cu(DDC)_2_. **B** and **C,** RAI uptake in parental 8505C cells (**B**) and human primary thyrocytes (**C**) following PAX8-siRNA depletion and Cu(DDC)_2_ treatment. CON – scrambled control siRNA. See also Supp Fig. S7A-S7D. **D,** RAI uptake in 8505C-NIS and TPC-1-NIS cells following NPL4- or VCP-siRNA depletion and Cu(DDC)_2_ treatment. CON – scrambled control siRNA. **E,** Western blot analysis of NIS, NPL4 and VCP protein in 8505C-NIS and TPC-1-NIS cells after NPL4- or VCP-siRNA depletion as described in **D**. **F** and **G,** Same as **D** and **E** but in human primary thyrocytes. **H,** NanoBiT evaluation of protein: protein interaction between NIS and VCP in live HeLa cells treated with CB5339 or Cu(DDC)_2_ at indicated doses. NanoBiT assay results at 20 min post-addition of Nano-Glo live cell assay substrate. See also Supp Fig. 7E. **I,** Schematic depicting NanoBRET assay to monitor proximity of NIS with highly abundant PM protein KRAS, as well as subcellular markers RAB5 (early endosome) and RAB11 (recycling endosome). Created with BioRender.com. **J,** NanoBRET evaluation of NIS PM (KRAS) and recycling endosome (RAB11) localisation in live HeLa cells treated with Cu(DDC)_2_. See also Supp Fig. S7F.

**Figure 5.**
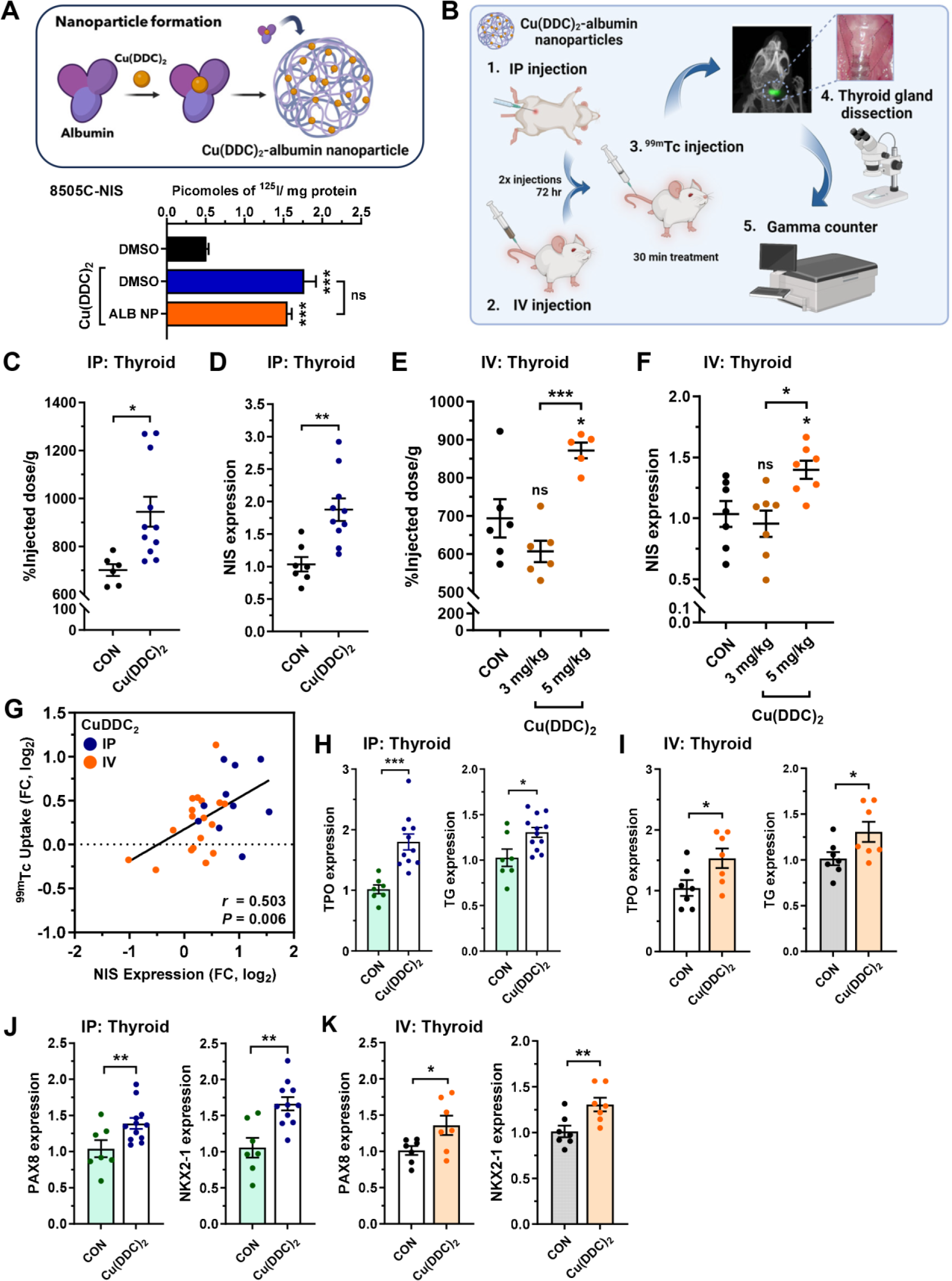
Targeting NIS function with Cu(DDC)_2_ enhances radionuclide uptake *in vivo*. **A,** (*above*) Schematic showing formation of Cu(DDC)_2_-albumin nanoparticles [Cu(DDC)_2_-ALB NP]. (*below*) RAI uptake in 8505C-NIS cells treated with Cu(DDC)_2_ in DMSO or encapsulated (ALB NP). Created in Biorender.com. See also Supp Fig. S8A. **B,** Schematic of steps used to examine the translatable potential of Cu(DDC)_2_ to enhance NIS function *in vivo*. Cu(DDC)_2_-ALB NP were given by either IP (1) or IV (2) injection prior to Technetium-99m pertechnetate (^99m^Tc) administration (3), thyroid gland dissection (4) and gamma counting (5). Created in Biorender.com. **C** and **D**, ^99m^Tc uptake (**C**; *n* = 6 - 11) and NIS mRNA levels (**D**) in thyroid glands dissected from WT BALB/c mice administered by IP injection with Cu(DDC)_2_-ALB NP. **E** and **F,** Same as **C** and **D** but Cu(DDC)_2_-ALB NP administered by IV injection at indicated doses (n = 5 – 7). **G,** Correlation analysis between thyroidal ^99m^Tc uptake (FC, log_2_) and NIS mRNA levels (FC, log_2_) in WT BALB/c mice injected with Cu(DDC)_2_ as outlined in **B**. **H,** Relative TPO and TG mRNA levels in thyroid glands dissected from WT BALB/c mice administered by IP injection with Cu(DDC)_2_-ALB NP. **I,** Same as **H** but Cu(DDC)_2_-ALB NP given by IV injection. **J,** Same as **H** but relative PAX8 and NKX2-1 mRNA levels. **K,** Same as **I** but relative PAX8 and NKX2-1 mRNA levels. Data presented as mean ± S.E.M.; ns, not significant; \**P* < 0.05; \*\**P* < 0.01; \*\*\**P* < 0.001.

**Figure 6.**
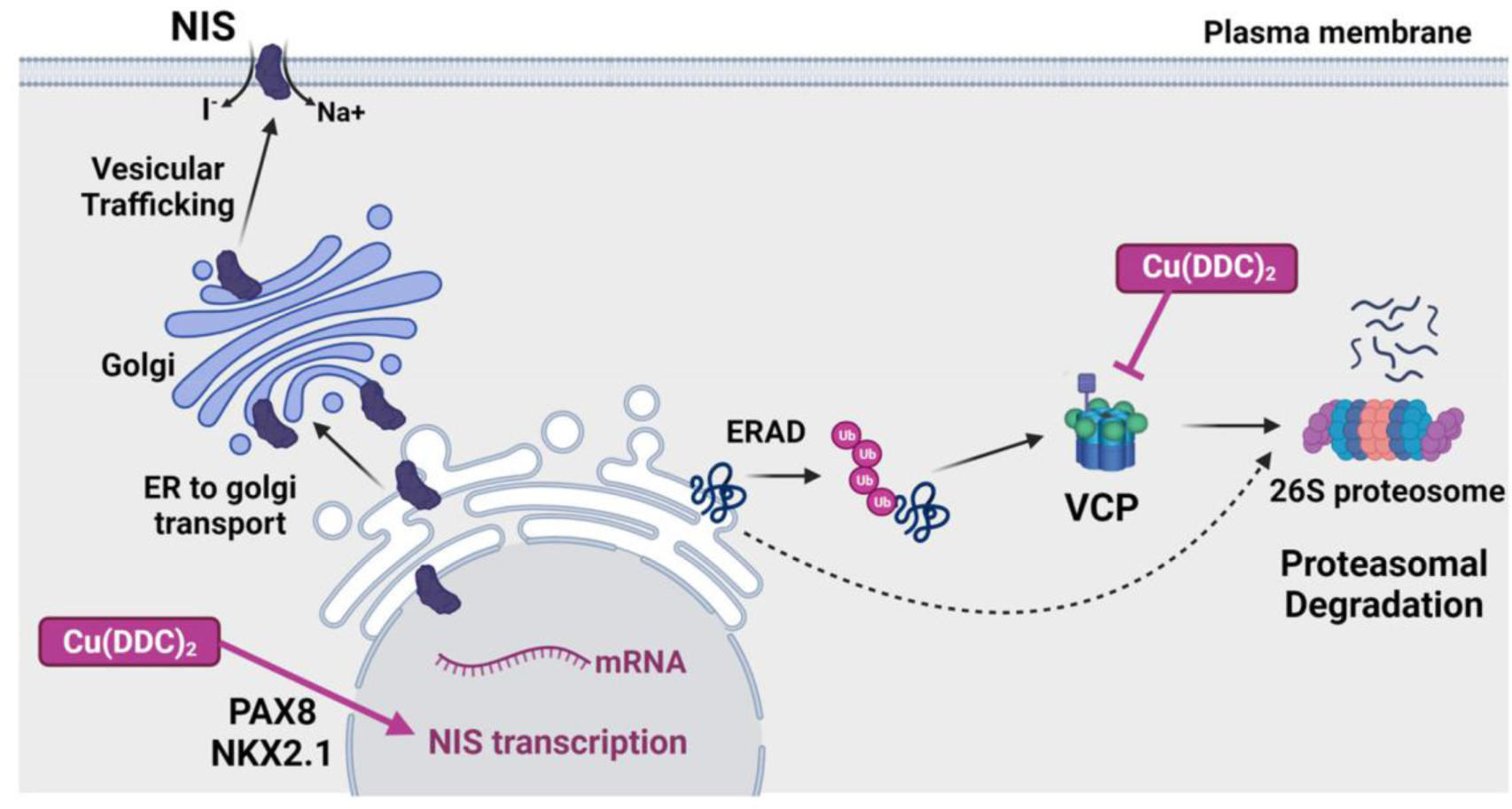
Schematic showing dual mechanistic impact of Cu(DDC)_2_ modulating NIS activity and RAI uptake in thyroid cells. (1) Cu(DDC)_2_ increases expression of NIS regulators such as PAX8 which enhance NIS transcription, and (2) Cu(DDC)_2_ does not require functional NPL4 expression in order to enhance RAI uptake, but does require VCP. Created with BioRender.com.

### Cu(DDC)_2_ induction of RAI uptake is dependent on PAX8 and VCP

To challenge the hypothesis that Cu(DDC)_2_ acts as a dual activator of NIS function, we first ablated PAX8, a master regulator of the thyroid differentiated phenotype [24], and examined whether Cu(DDC)_2_ retained its ability to stimulate RAI uptake. Critically, Cu(DDC)_2_ was unable to induce ^125^I uptake when PAX8 was depleted in thyroid cancer cells and primary thyrocytes (Fig. 4B and 4C; Supp Fig. 7A). In support of this, Cu(DDC)_2_ failed to induce NIS mRNA expression in primary thyrocytes when PAX8 was ablated (Supp Fig. S7B and S7C). Further, in contrast to PAX8, Cu(DDC)_2_ retained its induction of RAI uptake in 8505C-NIS and primary thyrocytes when CREM was depleted (Supp Fig. 7D).

Surprisingly, when we abrogated VCP and NPL4, Cu(DDC)_2_ retained activity in the absence of NPL4 but not VCP in thyroid cancer cells and primary thyrocytes (Fig. 4D-4G). Thus despite being a reported inhibitor of NPL4 [11], Cu(DDC)_2_ required functional VCP but not NPL4 expression in order to enhance RAI uptake. Subsequent NanoBiT assays in live HeLa cells revealed that, in contrast to CB5339, Cu(DDC)_2_ was unable to impact the stringency of the NIS: VCP interaction, and is therefore likely to be acting as a non-canonical VCP inhibitor (Fig. 4H; Supp Fig. S7E).

We next performed NanoBRET assays in live cells to determine whether Cu(DDC)_2_ can impact NIS subcellular localisation - the cardinal determinant of its symporter function - independently of transcriptional regulation. Utilising the plasma membrane (PM) marker KRAS [25], Cu(DDC)_2_ treatment was indeed associated with increased NIS localisation at the PM (Fig. 4J). We also found greater NIS in the endoplasmic reticulum (ER) to cis-golgi location (RAB1 marker; Supp Fig. S7F) but not in recycling endosomes (RAB11 marker; Fig. 4J). Collectively, these data are supportive that Cu(DDC)_2_ acts as a dual activator of NIS function, dependent on the distinct functionalities of PAX8 and VCP.

### Cu(DDC)_2_ enhances radionuclide uptake *in vivo*

Cu(DDC)_2_ is known to have poor *in vivo* solubility [26], limiting its potential clinical utility. Before elucidating the potency of Cu(DDC)_2_ *in vivo*, we nanoencapsulated the drug with albumin in an attempt to enhance its solubility and bioavailability (Fig. 5A). Importantly, nanoencapsulated Cu(DDC)_2_ retained the ability to stimulate RAI uptake in 8505C and TPC-1 cells in comparison to un-encapsulated drugs (Fig. 5A; Supp Fig. S8A), enabling us to progress to *in vivo* experiments as outlined (Fig. 5B).

In wild-type BALB/c mice, intraperitoneal (IP) administration of albumin nano-encapsulated Cu(DDC)_2_ significantly induced thyroidal uptake of technetium-99m (^99m^Tc) after 30 min compared to vehicle (∼40% increase; n = 6-11 per group; 3 mg/kg dose; *P* < 0.05; Fig. 5C). Thyroidal mRNA levels of NIS were also induced by 1.9-fold (*P* < 0.01; Fig. 5D). We next utilised intravenous (IV) injection to deliver doses of 3 and 5 mg/kg to BALB/c mice. Whereas no change was noted at 3 mg/kg compared to vehicle, the higher 5 mg/kg dose of nano-encapsulated Cu(DDC)_2_ significantly increased thyroidal ^99m^Tc-uptake (Fig. 5E), again accompanied by higher thyroidal NIS mRNA expression (Fig. 5F). No significant increases in ^99m^Tc uptake were detected in other major tissues (Supp Fig. S8B and S8C). Importantly, a significant positive correlation between thyroidal ^99m^Tc uptake and higher NIS mRNA levels (r_s_ = 0.503, *P* = 0.0169) was apparent in Cu(DDC)_2_-treated mice (Fig. 5G). Combined, our IP and IV administration data revealed that Cu(DDC)_2_ was able to significantly influence endogenous NIS activity *in vivo*.

We next investigated the mRNA expression of key thyroidal genes in response to drug treatment. Thyroid peroxidase (TPO) and thyroglobulin (TG) were both significantly induced in thyroid glands of mice treated with albumin nano-encapsulated Cu(DDC)_2_, whether delivered via IP or IV (Fig. 5H and 5I). Of particular significance, we further discovered that the key thyroid transcription factors PAX8 and NKX2-1 were elevated in Cu(DDC)_2_ treated mice compared to controls (Fig. 5J and 5K; Supp Fig. S8D-S8F), suggesting a mechanism for the upregulation of NIS expression in thyroid glands *in vivo*. Overall, we identify a new drug approach for inducing NIS expression and function *in vivo* and propose a model for a dual agonist function via the inhibition of VCP-mediated degradation coupled with the induction of NIS transcription (Fig. 6).

## Discussion

In addressing treatment failure in thyroid cancer, scant attention has been paid to targeting the mechanisms via which the sodium iodide symporter is trafficked and processed, with the majority of pre-clinical and clinical strategies being focussed around re-establishing the expression of NIS [27–29]. In attempting to modulate the intracellular processing of NIS we identified the proteostatic inhibitor disulfiram to be a potent enhancer of radioiodide uptake [7]. However, presumably due to unfavourable pharmacokinetics, disulfiram was incapable of increasing radionuclide uptake into murine thyroid glands. Unexpectedly, investigation of a copper metabolite of disulfiram led us to a compound which – rather than acting solely via disulfiram’s canonical proteostatic mechanism(s) – was a significant transcriptional regulator of NIS expression. Cu(DDC)_2_ treatment resulted in increased NIS localisation to the plasma membrane in living cells – as determined via NanoBRET – accompanied by greater transport activity *in vitro* and *in vivo*, and raised NIS mRNA levels.

In pre-clinical xenograft models of prostate cancer, disulfiram reduced tumour growth, but did so only when co-administered with copper [30]. In keeping with this, copper gluconate enhanced disulfiram’s impact upon radioiodide uptake in our assays, although copper itself was not able to stimulate ^125^I uptake. Multiple clinical trials in various neoplasias are currently appraising disulfiram in combination with copper gluconate [8]. However, there are concerns for adopting this approach, including potential mismatch of the pharmacokinetics of disulfiram and copper in systemic application, as well as a low dose of Cu(DDC)_2_ reaching the tumour site [31]. As previously mentioned, it has recently been shown that several of disulfiram’s anti-tumour properties are mediated via Cu(DDC)_2_ inhibition of the VCP cofactor NPL4 [11]. In contrast to the metal ion-dependent antitumor activity of disulfiram, direct delivery of Cu(DDC)_2_ may therefore reduce the dose of the drug needed and increase the therapeutic index [31].

VCP is a highly conserved multifunctional protein, with essential roles in pathways related to the ubiquitin-proteasome system, including ERAD, and is also central to non-proteolytic functions of ubiquitin signaling, including cell cycle regulation and Golgi biogenesis [32]. Cu(DDC)_2_ has been shown to segregate VCP from chromatin and into inactive agglomerates by disrupting NPL4 zinc finger motifs [11]. In our studies, we discovered a transcriptional impact of Cu(DDC)_2_ on NIS expression mediated via the key thyroid transcription factor PAX8: transient knockdown of PAX8 prevented NIS upregulation by Cu(DDC)_2_, as well as the subsequent induction of radioiodide uptake. However, whilst NPL4 was not obligate to this, the presence of VCP remained critical, and its expression was not altered by drug treatment. VCP has numerous cofactors in addition to NPL4, with up to 170 different post-translational modifications modulating their specific interactions [33]. Within this complexity we suggest that as well as transcriptional activation, Cu(DDC)_2_ impacts a facet of VCP biology which is independent of the cofactor NLP4.

As an alternative strategy to enhance radioiodide uptake *in vivo* by modulating NIS post-translational processing, we appraised the allosteric VCP inhibitor clotrimazole. This called upon our earlier finding that clotrimazole inhibits VCP-mediated ERAD of NIS [10], and our current data that the competitive ATPase VCPi CB5083 was ineffective in our preliminary mouse experiments. We constructed a total of 26 de novo compounds with improved predicted bioavailabilities, modelled on the clotrimazole structure in conjunction with docking studies of known VCP allosteric inhibitors. Of this pool, two compounds (C11 and C17) showed most promise in being effective at increasing radioiodide uptake and having improved logP values. Interestingly, we found that debulking the aryl groups in clotrimazole had the greatest overall impact on bioavailability correlating with enhanced NIS activity. This was presumably due to increased affinity of these compounds for the VCP binding pocket via removal of both bulky aryl groups. However, despite extensive efforts, the potency of our clotrimazole derivatives was over-shadowed by the disulfiram metabolite Cu(DDC)_2_, which ultimately proved effective *in vivo*.

Previous studies have shown that Cu(DDC)_2_ is poorly soluble [26]. Hence, in this study we nanoencapsulated Cu(DDC)_2_ in albumin to improve its *in vivo* stability and bioavailability, as reported by Skrott and colleagues [11]. Importantly, Cu(DDC)_2_-albumin retained *in vitro* impact upon NIS function, which emboldened us to progress to murine studies. Whether delivered via IV or IP routes, Cu(DDC)_2_-albumin stimulated thyroidal ^99m^Tc uptake in wild type BALB/c mice, implying a role in the normal physiological processing of NIS activity. Further studies will be needed to evaluate Cu(DDC)_2_-albumin nanoparticles in murine models of thyroid cancer, where we anticipate the effect may be significantly greater, given the transcriptional repression of NIS under such circumstances. Of particular note, Cu(DDC)_2_ was effective in parental 8505C, TPC-1, SW1736 and BCPAP thyroid cancer cells, all of which have markedly repressed expression of NIS and related transcription factors.

Mechanistically, our findings collectively suggest that Cu(DDC)_2_ acts as a dual activator of NIS function to enhance radioiodide uptake, given the known and potent impact of VCP on NIS protein degradation [10]. It is not, however, experimentally possible to compare the relative contributions of transcriptional activation versus NIS proteasomal degradation *in vivo*.

As drug treatments were short-term (72 hr), potential long-term side effects would presumably be obviated. Of note, BALB/c mice demonstrated no significant toxicity issues. We propose therefore that clinical trials are now merited, with patients given two IV injections of Cu(DDC)_2_-albumin nanoparticles, to appraise both drug dose and tolerability, and the impact on tracer radioiodide uptake into thyroid tumours.

In summary, we identify a new dual agonist of radioiodide uptake *in vivo*, with the potential to impact radioiodide therapy for patients with aggressive thyroid cancer who typically have poorer clinical outcomes.

## Acknowledgments

This work was supported by the Department of Defense (BC201532P1 to M.J. Campbell and C.J. McCabe). C.J. McCabe also received funding from the Medical Research Council (CiC/1001505) and British Thyroid Foundation (1002175). We acknowledge the labs of K. Pfleger and N. Lambert for kindly providing Venus-tagged constructs. We further acknowledge support from the Wellcome Trust and EPSRC funded Centre for Medical Engineering at King’s College London (203148/Z/16/Z), the Wellcome Multiuser Equipment Radioanalytical Facility (212885/Z/18/Z), and the EPSRC programme for Next Generation Molecular Imaging and Therapy with Radionuclides (EP/S019901/1).

## References

1. Schlumberger M, Brose M, Elisei R, Leboulleux S, Luster M, Pitoia F, et al. Definition and management of radioactive iodine-refractory differentiated thyroid cancer. Lancet Diabetes Endocrinol. 2014;2(5):356–8.

2. Sung H, Ferlay J, Siegel RL, Laversanne M, Soerjomataram I, Jemal A, et al. Global Cancer Statistics 2020: GLOBOCAN Estimates of Incidence and Mortality Worldwide for 36 Cancers in 185 Countries. CA Cancer J Clin. 2021;71(3):209–49.

3. Leboulleux S, Do Cao C, Zerdoud S, Attard M, Bournaud C, Lacroix L, et al. A Phase II Redifferentiation Trial with Dabrafenib-Trametinib and 131I in Metastatic Radioactive Iodine Refractory BRAF p.V600E-Mutated Differentiated Thyroid Cancer. Clin Cancer Res. 2023;29(13):2401–9.

4. Leboulleux S, Benisvy D, Taieb D, Attard M, Bournaud C, Terroir-Cassou-Mounat M, et al. MERAIODE: A Phase II Redifferentiation Trial with Trametinib and (131)I in Metastatic Radioactive Iodine Refractory RAS Mutated Differentiated Thyroid Cancer. Thyroid. 2023;33(9):1124–9.

5. Crispo F, Notarangelo T, Pietrafesa M, Lettini G, Storto G, Sgambato A, et al. BRAF Inhibitors in Thyroid Cancer: Clinical Impact, Mechanisms of Resistance and Future Perspectives. Cancers (Basel). 2019;11(9):1388.

6. Liu Y, Wang J, Hu X, Pan Z, Xu T, Xu J, et al. Radioiodine therapy in advanced differentiated thyroid cancer: Resistance and overcoming strategy. Drug Resist Updat. 2023;68:100939.

7. Read ML, Brookes K, Thornton CEM, Fletcher A, Nieto HR, Alshahrani M, et al. Targeting non-canonical pathways as a strategy to modulate the sodium iodide symporter. Cell Chem Biol. 2022;29(3):502–16 e7.

8. Kannappan V, Ali M, Small B, Rajendran G, Elzhenni S, Taj H, et al. Recent Advances in Repurposing Disulfiram and Disulfiram Derivatives as Copper-Dependent Anticancer Agents. Front Mol Biosci. 2021;8:741316.

9. Suh JJ, Pettinati HM, Kampman KM, O’Brien CP. The status of disulfiram: a half of a century later. J Clin Psychopharmacol. 2006;26(3):290–302.

10. Fletcher A, Read ML, Thornton CEM, Larner DP, Poole VL, Brookes K, et al. Targeting Novel Sodium Iodide Symporter Interactors ADP-Ribosylation Factor 4 and Valosin-Containing Protein Enhances Radioiodine Uptake. Cancer Res. 2020;80(1):102–15.

11. Skrott Z, Mistrik M, Andersen KK, Friis S, Majera D, Gursky J, et al. Alcohol-abuse drug disulfiram targets cancer via p97 segregase adaptor NPL4. Nature. 2017;552(7684):194-9.

12. Segura-Cabrera A, Tripathi R, Zhang X, Gui L, Chou TF, Komurov K. A structure- and chemical genomics-based approach for repositioning of drugs against VCP/p97 ATPase. Sci Rep. 2017;7:44912.

13. Pettersen EF, Goddard TD, C.C. H, G.S. C, Greenblatt DM, Meng EC, et al. UCSF Chimera--a visualization system for exploratory research and analysis. J Comput Chem 2004;25(13):1605–12.

14. Banerjee S, Bartesaghi A, Merk A, Rao P, Bulfer SL, Yan Y, et al. 2.3 A resolution cryo-EM structure of human p97 and mechanism of allosteric inhibition. Science. 2016;351(6275):871-5.

15. Tiulpakov A, White CW, Abhayawardana RS, See HB, Chan AS, Seeber RM, et al. Mutations of Vasopressin Receptor 2 Including Novel L312S Have Differential Effects on Trafficking. Mol Endocrinol. 2016;30(8):889–904.

16. Smith VE, Read ML, Turnell AS, Watkins RJ, Watkinson JC, Lewy GD, et al. A novel mechanism of sodium iodide symporter repression in differentiated thyroid cancer. Journal of Cell Science. 2009;122(Pt 18):3393–402.

17. Anderson DJ, Le Moigne R, Djakovic S, Kumar B, Rice J, Wong S, et al. Targeting the AAA ATPase p97 as an Approach to Treat Cancer through Disruption of Protein Homeostasis. Cancer Cell. 2015;28(5):653–65.

18. Roux B, Vaganay C, Vargas JD, Alexe G, Benaksas C, Pardieu B, et al. Targeting acute myeloid leukemia dependency on VCP-mediated DNA repair through a selective second-generation small-molecule inhibitor. Sci Transl Med. 2021;13(587):eabg1168.

19. Morrison BW, Doudican NA, Patel KR, Orlow SJ. Disulfiram induces copper-dependent stimulation of reactive oxygen species and activation of the extrinsic apoptotic pathway in melanoma. Melanoma Res. 2010;20(1):11–20.

20. Yip NC, Fombon IS, Liu P, Brown S, Kannappan V, Armesilla AL, et al. Disulfiram modulated ROS-MAPK and NFkappaB pathways and targeted breast cancer cells with cancer stem cell-like properties. Br J Cancer. 2011;104(10):1564–74.

21. Chai W, Ye F, Zeng L, Li Y, Yang L. HMGB1-mediated autophagy regulates sodium/iodide symporter protein degradation in thyroid cancer cells. J Exp Clin Cancer Res. 2019;38(1):325.

22. Lambert SA, Jolma A, Campitelli LF, Das PK, Yin Y, Albu M, et al. The Human Transcription Factors. Cell. 2018;172(4):650–65.

23. Riesco-Eizaguirre G, Santisteban P, De la Vieja A. The complex regulation of NIS expression and activity in thyroid and extrathyroidal tissues. Endocr Relat Cancer. 2021;28(10):T141–T65.

24. Pasca di Magliano M, Di Lauro R, Zannini M. Pax8 has a key role in thyroid cell differentiation. Proc Natl Acad Sci U S A. 2000;97(24):13144–9.

25. Lan TH, Liu Q, Li C, Wu G, Lambert NA. Sensitive and high resolution localization and tracking of membrane proteins in live cells with BRET. Traffic. 2012;13(11):1450–6.

26. Skrott Z, Cvek B. Diethyldithiocarbamate complex with copper: the mechanism of action in cancer cells. Mini Rev Med Chem. 2012;12(12):1184–92.

27. Chakravarty D, Santos E, Ryder M, Knauf JA, Liao XH, West BL, et al. Small-molecule MAPK inhibitors restore radioiodine incorporation in mouse thyroid cancers with conditional BRAF activation. J Clin Invest. 2011;121(12):4700–11.

28. Ullmann TM, Liang H, Moore MD, Al-Jamed I, Gray KD, Limberg J, et al. Dual inhibition of BRAF and MEK increases expression of sodium iodide symporter in patient-derived papillary thyroid cancer cells in vitro. Surgery. 2020;167(1):56–63.

29. Yu Q, Zhang X, Li L, Zhang C, Huang J, Huang W. Molecular basis and targeted therapies for radioiodine refractory thyroid cancer. Asia Pac J Clin Oncol. 2023;19(3):279–89.

30. Safi R, Nelson ER, Chitneni SK, Franz KJ, George DJ, Zalutsky MR, et al. Copper signaling axis as a target for prostate cancer therapeutics. Cancer Res. 2014;74(20):5819–31.

31. Kang X, Jadhav S, Annaji M, Huang CH, Amin R, Shen J, et al. Advancing Cancer Therapy with Copper/Disulfiram Nanomedicines and Drug Delivery Systems. Pharmaceutics. 2023;15(6). 1567.

32. Braxton JR, Southworth DR. Structural insights of the p97/VCP AAA+ ATPase: How adapter interactions coordinate diverse cellular functionality. J Biol Chem. 2023;299(11):105182.

33. Hanzelmann P, Schindelin H. The Interplay of Cofactor Interactions and Post-translational Modifications in the Regulation of the AAA+ ATPase p97. Front Mol Biosci. 2017;4:21.

