## Supplementary Figures for "Dual agonism of sodium iodide symporter function *in vivo*"

### SUPPLEMENTARY FIGURE 1

## A 8505C

**Laboratory Report**

Test Requested: Cell Line Authentication
Case Number: C-30880-1
Date Sample Received: 14/02/2024
Date Sample Tested: 14/02/2024
Date Sample Reported: 19/02/2024

| Sample Name / Cell Line | Cell Line Source / Profile Source | Sample Number | DNA Number |
|---|---|---|---|
| 8505C | University of Birmingham | S-1070452 | D-1070452 |
| 8505C (CVCL_1054) | Cellosaurus Database | N/A | N/A |

**Table of Allelic Data**

| | Genotypes | | |
|---|---|---|---|
| STR Locus | 8505C (Test Sample) | 8505C (CVCL_1054) (Comparison Sample) | Match vs. Mis-Match |
| D5S818 | 10 \| 11 | 10 \| 11 | Match |
| D13S317 | 13 \| 13 | 13 \| 13 | Match |
| D7S820 | 10 \| 10 | 10 \| 10 | Match |
| D16S539 | 12 \| 12 | 12 \| 12 | Match |
| vWA | 17 \| 19 | 17 \| 19 | Match |
| TH01 | 6 \| 9 | 6 \| 9 | Match |
| TPOX | 10 \| 11 | 10 \| 11 | Match |
| CSF1PO | 12 \| 13 | 12 \| 13 | Match |
| AMEL | X \| X | X \| X | Match |
| D3S1358 | 16 \| 17 | 16 \| 17 | Match |
| D21S11 | 28 \| 32.2 | 28 \| 32.2 | Match |
| D18S51 | 16 \| 16 | 16 \| 16 | Match |
| Penta E | 12 \| 15 | 12 \| 15 | Match |
| Penta D | 9 \| 10 | 9 \| 10 | Match |
| D8S1179 | 10 \| 13 | 10 \| 13 | Match |
| FGA | 23 \| 23 | 23 \| 23 | Match |
| D19S433 | 13 \| 14 | 13 \| 14 | Match |
| D2S1338 | 17 \| 24 | 17 \| 24 | Match |

Matching Percentage: **100%**
Outcome: **Related**

#### B TPC-1

**Laboratory Report**

Test Requested: Cell Line Authentication
Case Number: C-30880-2
Date Sample Received: 14/02/2024
Date Sample Tested: 14/02/2024
Date Sample Reported: 19/02/2024

| Sample Name / Cell Line | Cell Line Source / Profile Source | Sample Number | DNA Number |
|---|---|---|---|
| TPC1 | University of Birmingham | S-1070453 | D-1070453 |
| TPC-1 (CVCL_6298) | Cellosaurus Database | N/A | N/A |

**Table of Allelic Data**

| | Genotypes | | |
|---|---|---|---|
| STR Locus | TPC1 (Test Sample) | TPC-1 (CVCL_6298) (Comparison Sample) | Match vs. Mis-Match |
| D5S818 | 8 \| 10 | 8 \| 10 | Match |
| D13S317 | 11 \| 12 | 11 \| 12 | Match |
| D7S820 | 11 \| 11 | 11 \| 11 | Match |
| D16S539 | 9 \| 9 | 9 \| 9 | Match |
| vWA | 14 \| 18 | 14 \| 18 | Match |
| TH01 | 9 \| 9 | 9 \| 9 | Match |
| TPOX | 11 \| 11 | 11 \| 11 | Match |
| CSF1PO | 11 \| 12 | 11 \| 12 | Match |
| AMEL | X \| X | X \| X | Match |
| D3S1358 | 16 \| 17 | 16 \| 17 | Match |
| D21S11 | 30 \| 31.2 | 30 \| 31.2 | Match |
| D18S51 | 13 \| 16 | 13 \| 16 | Match |
| Penta E | 18 \| 18 | 18 \| 18 | Match |
| Penta D | 9 \| 13 | 9 \| 13 | Match |
| D8S1179 | 11 \| 17 | 11 \| 17 | Match |
| FGA | 20 \| 21 | 20 \| 21 | Match |
| D19S433 | 13 \| 13 | ǂ \| ǂ | Mis-Match |
| D2S1338 | 16 \| 23 | ǂ \| ǂ | Mis-Match |

Matching Percentage: **100%**
Outcome: **Related**

## C SW1736

**Laboratory Report**

Test Requested: Cell Line Authentication
Case Number: C-30880-4
Date Sample Received: 14/02/2024
Date Sample Tested: 14/02/2024
Date Sample Reported: 19/02/2024

| Sample Name / Cell Line | Cell Line Source / Profile Source | Sample Number | DNA Number |
|---|---|---|---|
| SW1736 | University of Birmingham | S-1070455 | D-1070455 |
| SW1736 (CVCL_3883) | Cellosaurus Database | N/A | N/A |

**Table of Allelic Data**

| | Genotypes | | |
|---|---|---|---|
| STR Locus | SW1736 (Test Sample) | SW1736 (CVCL_3883) (Comparison Sample) | Match vs. Mis-Match |
| D5S818 | 12 \| 13 | 12 \| 13 | Match |
| D13S317 | 11 \| 12 | 11 \| 12 | Match |
| D7S820 | 8 \| 11 | 8 \| 11 | Match |
| D16S539 | 11 \| 12 | 11 \| 12 | Match |
| vWA | 16 \| 19 | 16 \| 19 | Match |
| TH01 | 6 \| 6 | 6 \| 6 | Match |
| TPOX | 11 \| 11 | 11 \| 11 | Match |
| CSF1PO | 12 \| 12 | 12 \| 12 | Match |
| AMEL | X \| X | X \| X | Match |
| D3S1358 | 16 \| 17 | 16 \| 17 | Match |
| D21S11 | 29 \| 31 | 29 \| 31 | Match |
| D18S51 | 14 \| 14 | 14 \| 14 | Match |
| Penta E | 11 \| 17 | 11 \| 17 | Match |
| Penta D | 12 \| 12 | 12 \| 12 | Match |
| D8S1179 | 13 \| 14 | 13 \| 13 | Mis-Match |
| FGA | 22 \| 22 | 22 \| 22 | Match |
| D19S433 | 14 \| 14 | ǂ \| ǂ | Mis-Match |
| D2S1338 | 19 \| 25 | ǂ \| ǂ | Mis-Match |

Matching Percentage: **98%**
Outcome: **Related**

#### D BCPAP

**Laboratory Report**

Test Requested: Cell Line Authentication
Case Number: C-30880-5
Date Sample Received: 14/02/2024
Date Sample Tested: 14/02/2024
Date Sample Reported: 19/02/2024

| Sample Name / Cell Line | Cell Line Source / Profile Source | Sample Number | DNA Number |
|---|---|---|---|
| BCPAP | University of Birmingham | S-1070456 | D-1070456 |
| B-CPAP (CVCL_0153) | Cellosaurus Database | N/A | N/A |

**Table of Allelic Data**

| | Genotypes | | |
|---|---|---|---|
| STR Locus | BCPAP (Test Sample) | B-CPAP (CVCL_0153) (Comparison Sample) | Match vs. Mis-Match |
| D5S818 | 10 \| 11 | 10 \| 11 | Match |
| D13S317 | 12 \| 12 | 12 \| 12 | Match |
| D7S820 | 10 \| 10 | 10 \| 10 | Match |
| D16S539 | 11 \| 12 | 11 \| 12 | Match |
| vWA | 14 \| 17 | 14 \| 17 | Match |
| TH01 | 6 \| 9.3 | 6 \| 9.3 | Match |
| TPOX | 8 \| 11 | 8 \| 11 | Match |
| CSF1PO | 13 \| 13 | 13 \| 13 | Match |
| AMEL | X \| X | X \| X | Match |
| D3S1358 | 16 \| 17 | 16 \| 17 | Match |
| D21S11 | 30 \| 31.2 | 30 \| 30 | Mis-Match |
| D18S51 | 13 \| 17 | 13 \| 17 | Match |
| Penta E | 5 \| 12 | 5 \| 12 | Match |
| Penta D | 10 \| 11 | 10 \| 11 | Match |
| D8S1179 | 12 \| 13 | 12 \| 13 | Match |
| FGA | 20 \| 23 | 20 \| 23 | Match |
| D19S433 | 13.2 \| 15 | 14 \| 15 | Mis-Match |
| D2S1338 | 18 \| 18 | 18 \| 18 | Match |

Matching Percentage: **95%**
Outcome: **Related**

**Figure S1.** Representative STR profiles of cell lines. Examples of cell line authentication reports for **A**, 8505C, **B**, TPC-1, **C**, SW1736 and **D**, BCPAP cells generated by NorthGene (Biofortuna). DNA profiles were compared to profiles located on the Cellosaurus database. Analysis was conducted using 8 loci allowing identical matches and approximate power of discrimination of 1 in 1,000,000,000. Cell lines were considered to be related with a matching percentage > 80%.

#### SUPPLEMENTARY FIGURE 2

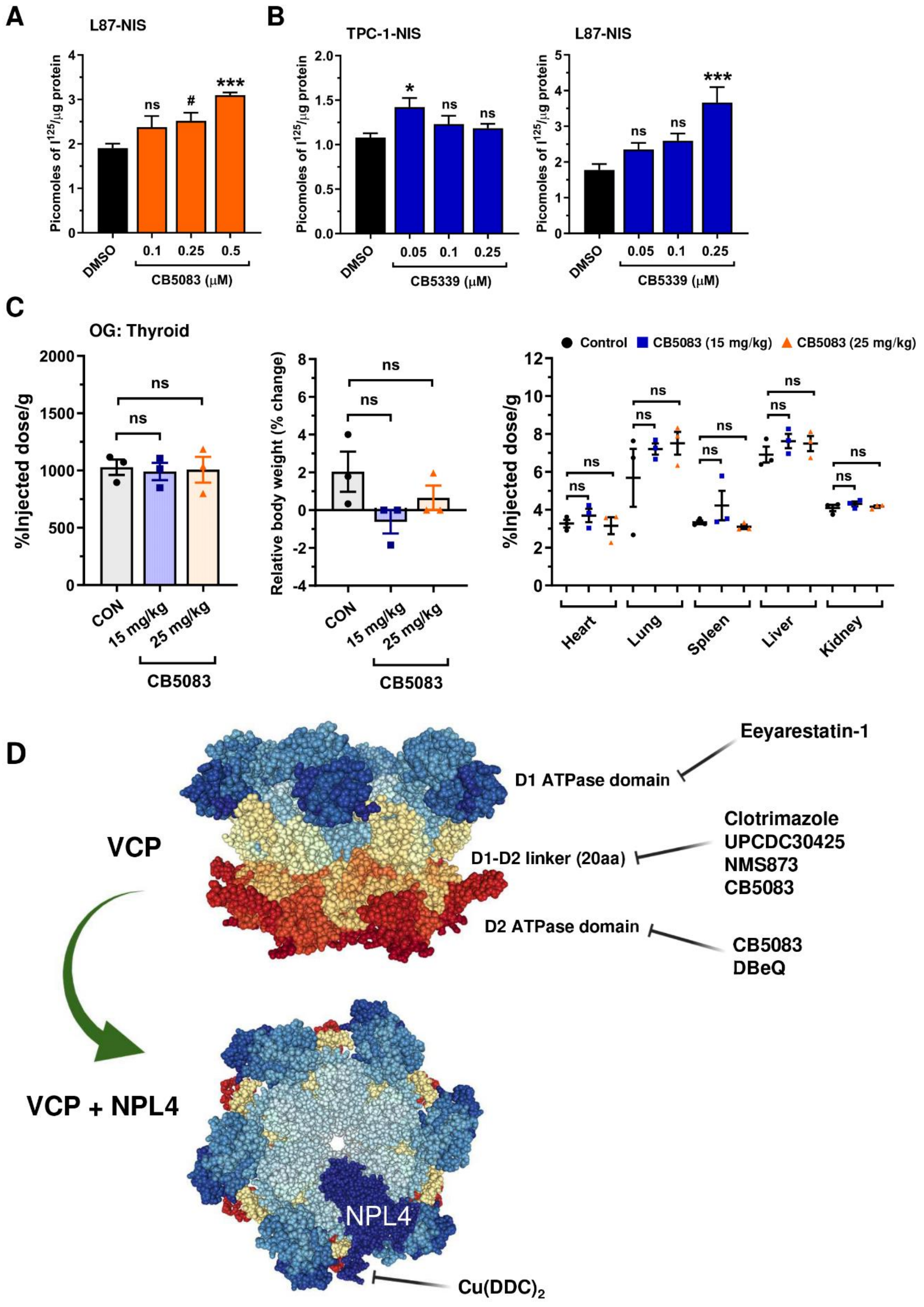

**Figure S2.** VCP inhibitors CB5083 and CB5339 enhance RAI uptake *in vitro*. **A,** RAI uptake in L87-NIS cells treated with CB5083 at indicated doses for 24 hr. **B,** RAI uptake in TPC-1-NIS and L87-NIS cells treated with CB5339 at indicated doses for 24 hr. **C,** $^{99m}$Tc uptake ($n$ = 3) in thyroid glands dissected from wild-type (WT) BALB/c mice administered by oral gavage with CB5083 at indicated doses. Mice were given a total of 4 daily doses. (*middle*) Body weight change (%) in WT BALB/c mice administered with CB5083 at indicated doses versus controls ($n$ = 3 per group). (*right*) Distribution of $^{99m}$Tc uptake across the indicated tissues harvested from WT BALB/c mice administered with CB5083 at indicated doses versus controls ($n$ = 3 per group). **D,** Schematic of VCP structure and regions targeted by VCP inhibitors. Targeted inhibition of the D1 to D2 20 residue linker region prevents conformational changes required for VCP function, while targeting the D2 domain inhibits ATPase enzyme activity. (*lower*) NPL4 binds to the N-domain of VCP. $Cu(DDC)_2$ binds NPL4 and induces its aggregation, which disables the p97–NPL4–UFD1 pathway (11). Data presented as mean ± S.E.M., one-way ANOVA followed by Dunnett's or Tukey's post hoc test (ns, not significant; $^*P < 0.05$; $^{***}P < 0.001$), or unpaired two-tailed t-test ($^{\#}P < 0.05$).

### SUPPLEMENTARY FIGURE 3

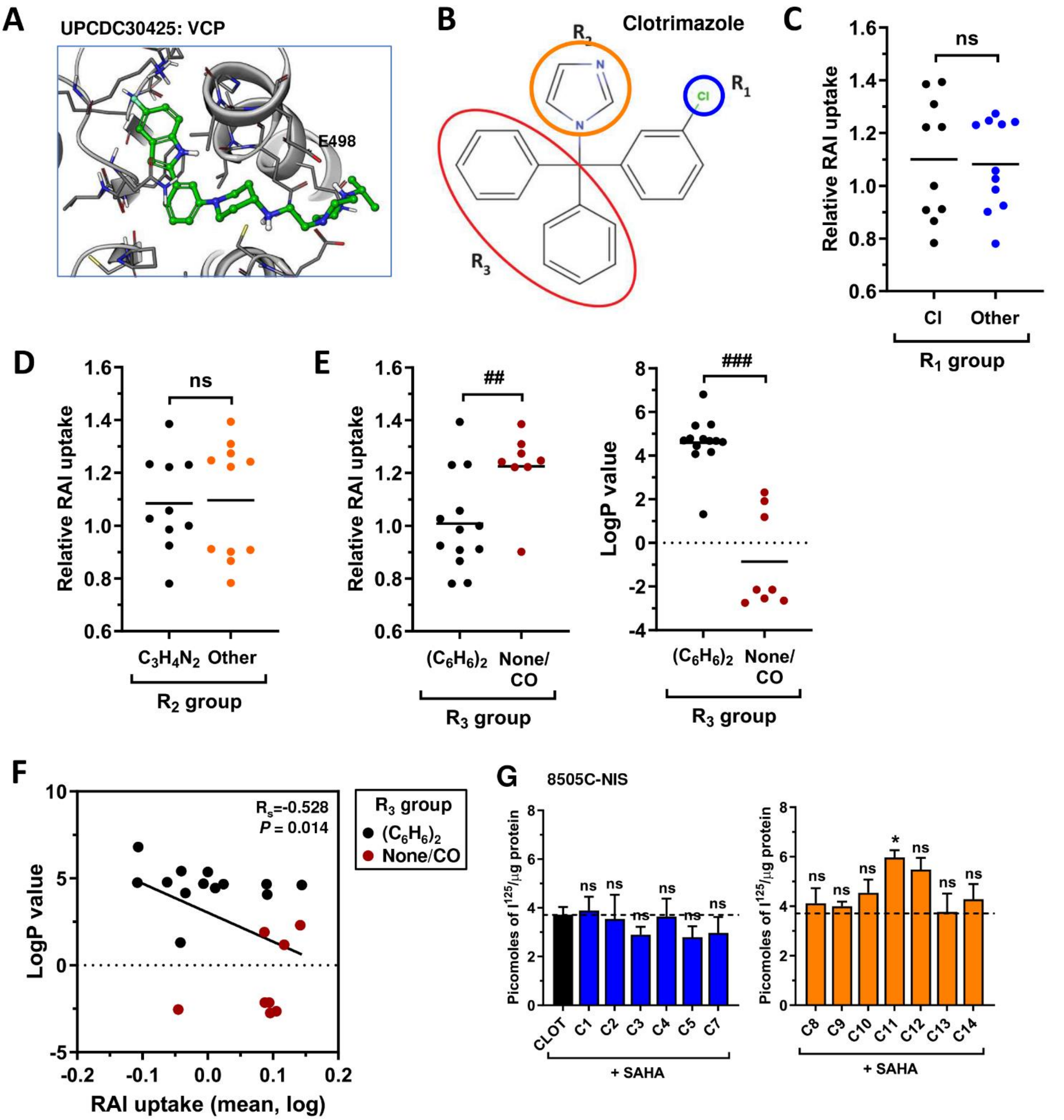

**Figure S3.** Debulking of phenyl groups increases the efficacy of clotrimazole analogues to enhance RAI uptake. **A,** Modelling the binding of UPCDC30425 to VCP. **B,** Chemical structure of clotrimazole highlighting modifications at the chloro-substituted aryl ring ($R_1$), imidazole ring ($R_2$) and two aryl substituent groups ($R_3$). **C,** Mean RAI uptake in thyroid cancer cell lines (8505C-NIS, TPC-1-NIS) treated with clotrimazole analogues with an intact ($R_1$ group-Cl) or modified chloro-substituted aryl ring ($R_1$ group-Other). See Supp Table S1 for details of clotrimazole derivatives. **D,** Same as **C** but comparing clotrimazole analogues with an intact ($R_2$ group-$C_3H_4N_2$) or modified imidazole ring ($R_2$ group-Other). **E,** (*left*) Same as **C** but comparing clotrimazole analogues with an intact [$R_3$ group-($C_6H_6)_2$] or modified aryl substituent groups ($R_3$ group-None/CO). (*right*) Comparison of logP values of clotrimazole analogues with an intact or modified $R_3$ group. **F,** Correlation analysis between mean RAI uptake and logP values of clotrimazole analogues with an intact or modified $R_3$ group. **G,** RAI uptake in 8505C-NIS cells treated with 21 compounds (C1-C21; 12 hr) in combination with SAHA (24 hr) versus clotrimazole (CLOT, C6) + SAHA. Dashed line represents normalised RAI uptake using CLOT in combination with SAHA. Data presented as mean ± S.E.M., one-way ANOVA followed by Dunnett's post hoc test (ns, not significant; $^{*}P < 0.05$), or unpaired two-tailed t-test ($^{\#\#}P < 0.01$; $^{\#\#\#}P < 0.001$).

#### SUPPLEMENTARY FIGURE 4

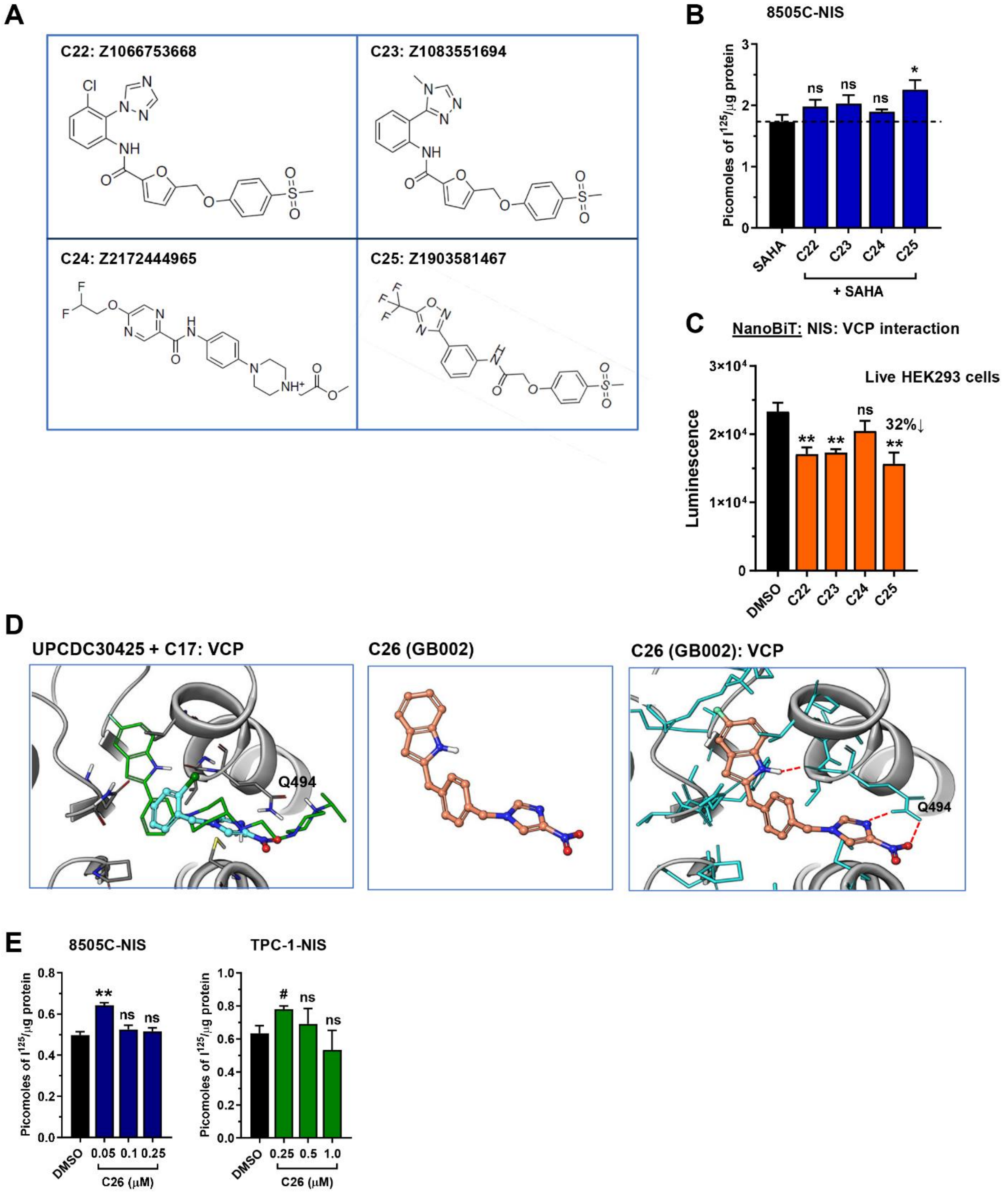

**Figure S4.** Rational drug design of VCP inhibitors to enhance NIS function. **A,** Chemical structures of compounds C22-C25. **B,** RAI uptake in 8505C-NIS cells treated with 4 compounds (C22-C25) in combination with SAHA versus SAHA alone. Dashed line represents RAI uptake using SAHA alone. **C,** NanoBiT evaluation of protein: protein interaction between NIS and VCP in live HEK293 cells treated with compounds C22-C25. NanoBiT assay results at 20 min post-addition of Nano-Glo live cell assay substrate. **D,** (*left*) Modelling of UPCDC30425 in the VCP allosteric binding pocket showed that the piperazine tail extended farther into the binding pocket than clotrimazole analogue C17, as evidenced by overlapping UPCDC30425 (green) with compound C17 (cyan). Based on this, GB002 was proposed as a modified compound C17 mimicking the piperazine tail of UPCDC30425 to capture further interactions with VCP residues forming the binding pocket. (*middle*) Chemical structure of compound C26 (GB002). (*right*) GB002 modelled in the binding pocket of VCP. Dashed lines represent predicted hydrogen bonds with residues (e.g. Q494) in the allosteric binding pocket of VCP. **E,** RAI uptake of 8505C-NIS and TPC-1-NIS cells treated with compound C26 (GB002) at indicated doses for 24 hr. Data presented as mean ± S.E.M., one-way ANOVA followed by Dunnett's post hoc test (ns, not significant; $^{*}P < 0.05$; $^{**}P < 0.01$), or unpaired two-tailed t-test ($^{\#}P < 0.05$).

#### SUPPLEMENTARY FIGURE 5

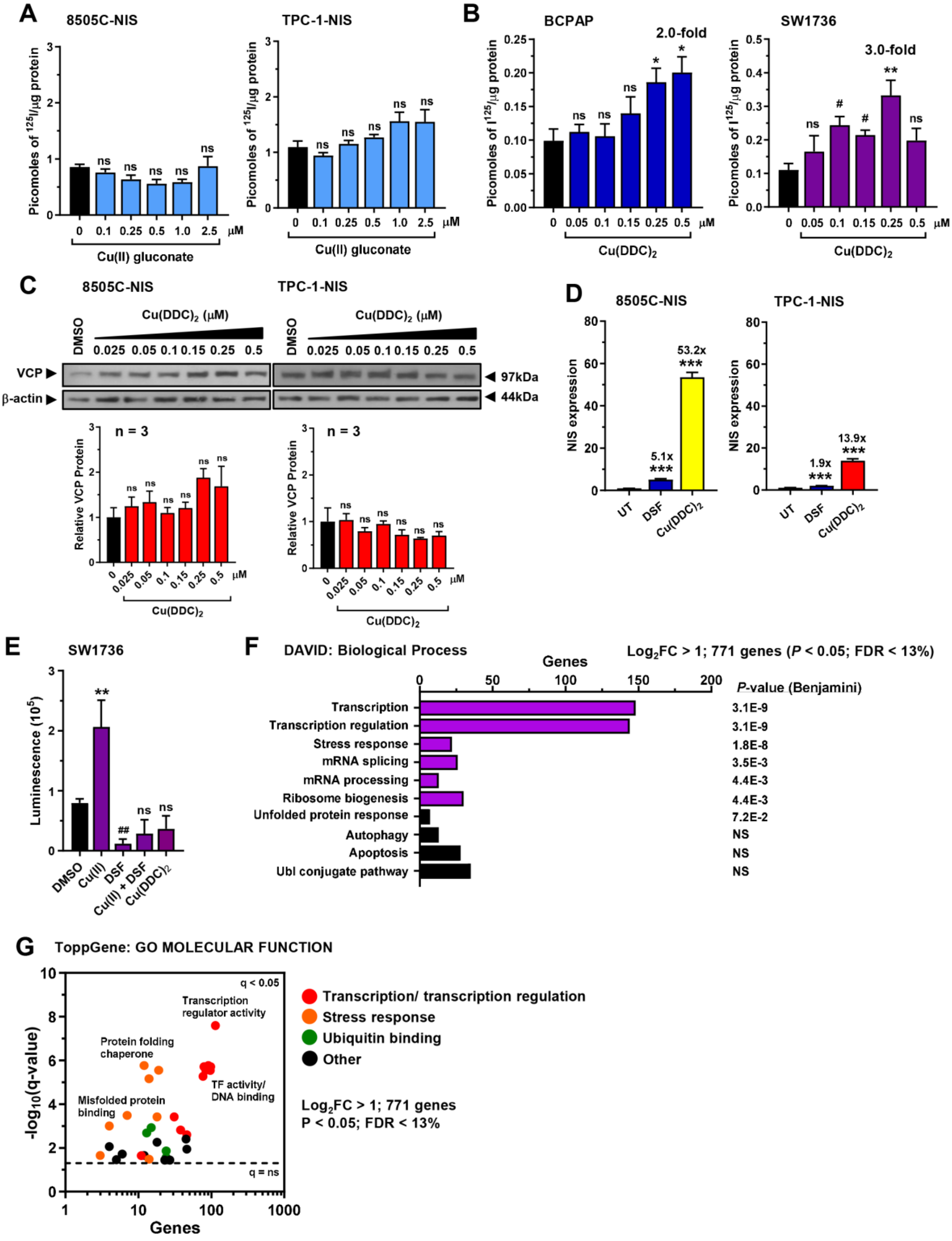

**Figure S5.** Cu(DDC)$_2$ elicits a transcriptional response to induce NIS function. **A,** RAI uptake of 8505C-NIS and TPC-1-NIS cells treated with copper gluconate [Cu(II)] at indicated doses for 24 hr. **B,** RAI uptake of BCPAP and SW1736 cells treated with Cu(DDC)$_2$ at indicated doses for 24 hr. **C,** Western blot analysis of VCP expression in 8505C-NIS and TPC-1-NIS cells treated with Cu(DDC)$_2$ at indicated doses for 24 hr. (*below*) Quantification of VCP protein levels ($n$ = 3). **D,** Relative NIS mRNA levels in 8505C-NIS and TPC-1-NIS cells treated with either DSF or Cu(DDC)$_2$. **E,** ROS-Glo assay evaluation of $H_2O_2$ levels in SW1736 cells treated with Cu(II), DSF and Cu(DDC)$_2$ versus DMSO as indicated. **F,** DAVID functional classification of top 771 differentially expressed genes ($\log_2$FC > 1, $P < 0.05$, FDR < 13%) in parental 8505C cells treated with 0.25 μM Cu(DDC)2 versus UT. **G,** Same as **F** but ToppGene used to classify the top 771 differentially expressed genes ($\log_2$FC > 1, $P < 0.05$, FDR < 13%). Data presented as mean ± S.E.M., one-way ANOVA followed by Dunnett's post hoc test (ns, not significant; $^*P < 0.05$; $^{**}P < 0.01$; $^{***}P < 0.001$), or unpaired two-tailed t-test ($^{\#}P < 0.05$; $^{\#\#}P < 0.01$).

#### SUPPLEMENTARY FIGURE 6

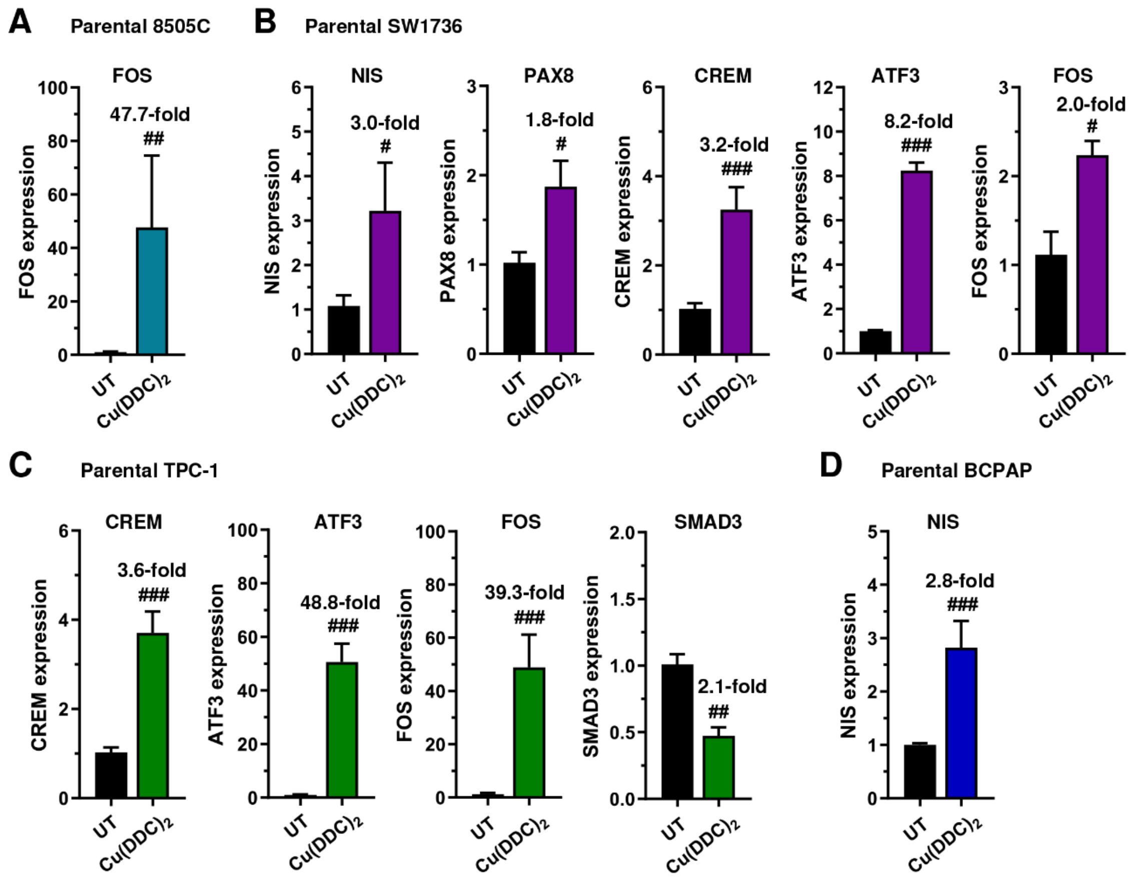


**Figure S6.** Cu(DDC)$_2$ induces expression of transcription factors in thyroid cells. **A,** Relative FOS mRNA levels in parental 8505C cells treated with Cu(DDC)$_2$. **B,** Relative NIS, PAX8, CREM, ATF3 and FOS mRNA levels in parental SW1736 cells treated with Cu(DDC)$_2$. **C,** Relative CREM, ATF3, FOS and SMAD3 mRNA levels in parental TPC-1 cells treated with Cu(DDC)$_2$. **D,** Relative NIS mRNA levels in parental BCPAP cells treated with Cu(DDC)$_2$. Data presented as mean ± S.E.M., unpaired two-tailed t-test ($^{\#}P < 0.05$; $^{\#\#}P < 0.01$; $^{\#\#\#}P < 0.001$).

### SUPPLEMENTARY FIGURE 7

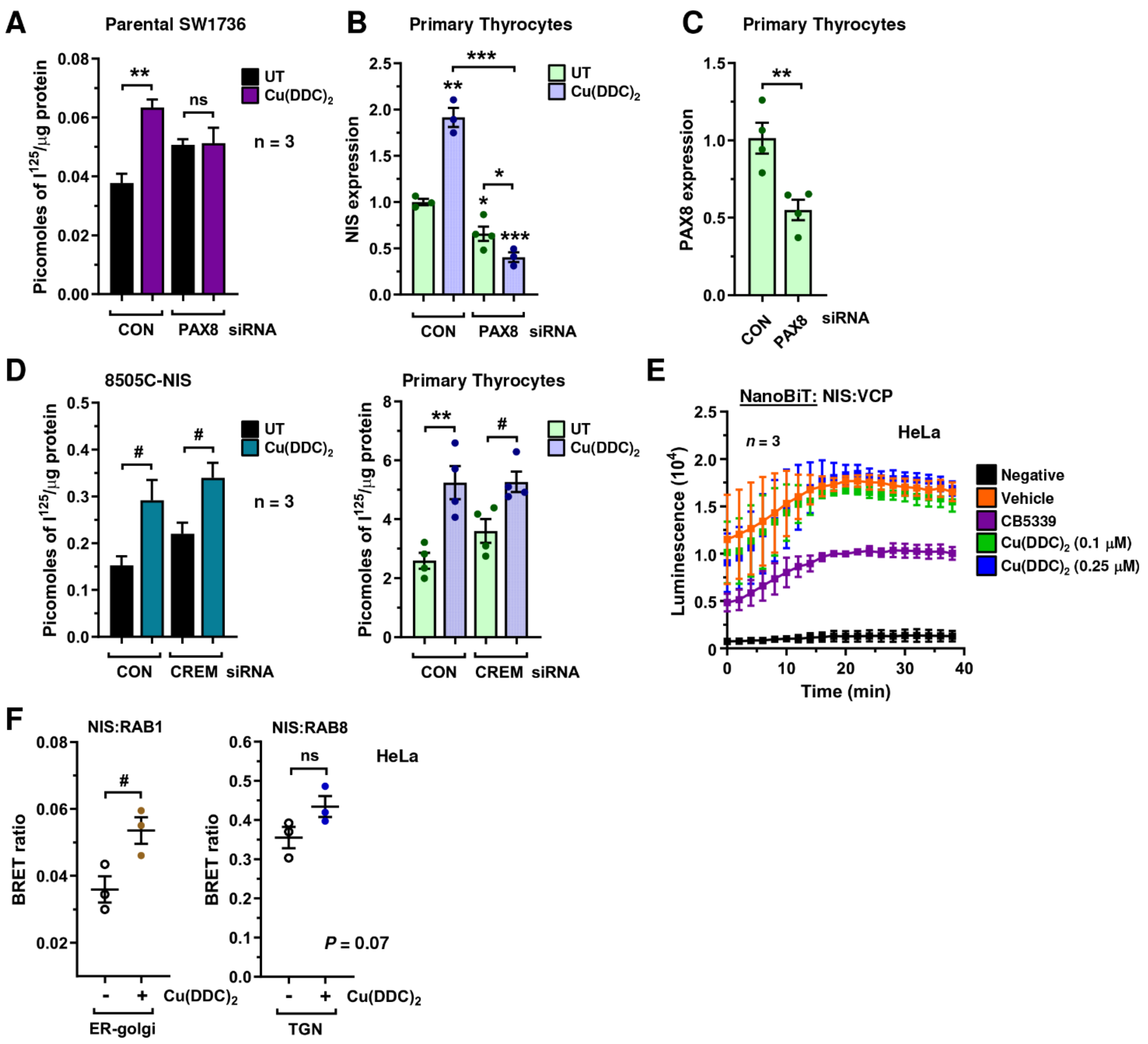

**Figure S7.** Cu(DDC)$_2$ induction of NIS mRNA and RAI uptake is dependent on PAX8. **A,** RAI uptake in parental SW1736 cells following PAX8-siRNA depletion and Cu(DDC)$_2$ treatment. CON – scrambled control siRNA. **B,** Relative NIS mRNA levels in human primary thyrocytes following PAX8-siRNA depletion and Cu(DDC)$_2$ treatment. **C,** Relative PAX8 mRNA levels in human primary thyrocytes following PAX8-siRNA depletion. **D,** RAI uptake in parental 8505C cells and human primary thyrocytes following CREM-siRNA depletion and Cu(DDC)$_2$ treatment. CON – scrambled control siRNA. **E,** NanoBiT evaluation of protein: protein interaction between NIS and VCP in living HeLa cells treated with CB5339 or Cu(DDC)$_2$ at indicated doses ($n$ = 3). **F,** NanoBRET evaluation of NIS ER-golgi (RAB1) and trans-golgi network (RAB8) localisation in live HeLa cells treated with Cu(DDC)$_2$ ($n$ = 3). Data presented as mean ± S.E.M., one-way ANOVA followed by Dunnett's or Tukey's post hoc test (ns, not significant; $*P < 0.05$; $**P < 0.01$; $***P < 0.001$), or unpaired two-tailed t-test ($^{\#}P < 0.05$).

#### SUPPLEMENTARY FIGURE 8

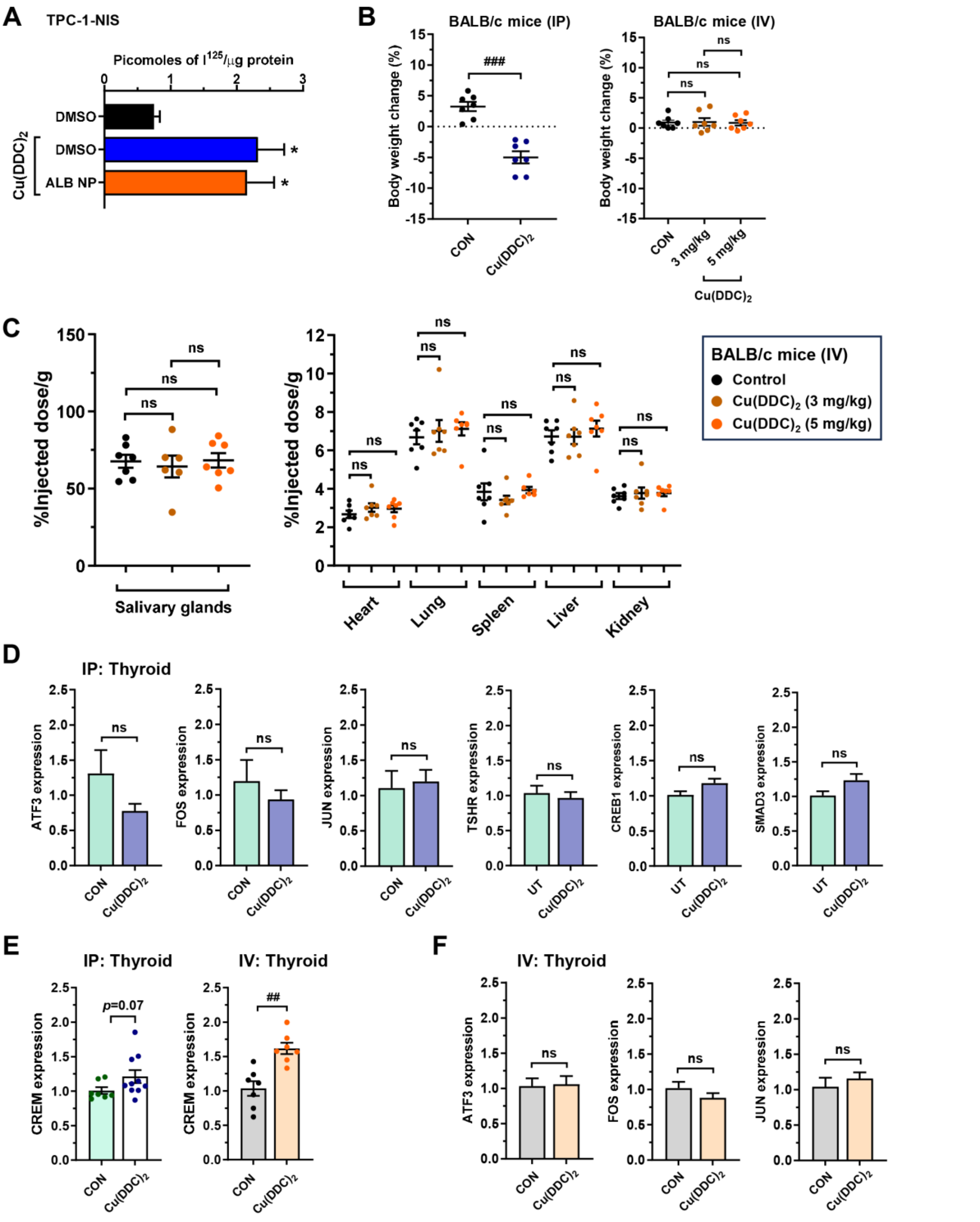


271

272

273

274

**Figure S8.** Biological effects of $Cu(DDC)_2$ and validatory experiments *in vivo*. **A,** RAI uptake in TPC-1-NIS cells treated with $Cu(DDC)_2$ in DMSO or encapsulated in albumin nanoparticles (ALB NP). **B,** Body weight change (%) in wild-type (WT) BALB/c mice administered with $Cu(DDC)_2$-ALB NP at indicated doses via IP (*left*) or IV injection (*right*) versus controls ($n$ = 5-7 per group). **C,** Distribution of $^{99m}Tc$ uptake across the indicated tissues harvested from WT BALB/c mice administered with $Cu(DDC)_2$-ALB NP at indicated doses versus controls ($n$ = 5-7 per group). **D,** Relative ATF3, FOS, JUN, TSHR, CREB1 and SMAD3 mRNA levels in thyroid glands dissected from WT BALB/c mice administered by IP injection with $Cu(DDC)_2$-ALB NP. **E,** Relative CREM mRNA levels in thyroid glands dissected from WT BALB/c mice administered by IP (*left*) or IV injection (*right*) with $Cu(DDC)_2$-ALB NP. **F,** Same as **D** but relative thyroidal ATF3, FOS and JUN mRNA levels in WT BALB/c mice administered by IV injection with $Cu(DDC)_2$-ALB NP. Data presented as mean ± S.E.M., one-way ANOVA followed by Dunnett's or Tukey's post hoc test (ns, not significant; $^{*}P < 0.05$), or unpaired two-tailed t-test (ns, not significant; $^{\#\#}P < 0.01$; $^{\#\#\#}P < 0.001$).

### SUPPLEMENTARY TABLE 1

| Clotrimazole derivatives | Modifications[1] | | | LogP | RAI (v SAHA + C6)[2] | | | RAI (v C6)[3] |
|---|---|---|---|---|---|---|---|---|
| | $R_1$ | $R_2$ | $R_3$ | | TPC-1-NIS | 8505C-NIS | Mean value | Mean value |
| CLOT/ C6 | Cl | $C_3H_4N_2$ | $(C_6H_6)_2$ | 5.37 | 1.00 | 1.00 | 1.00 | 1.00 |
| C1 | COOH | $C_3H_4N_2$ | $(C_6H_6)_2$ | 4.07 | 1.42 | 1.05 | 1.23 | 1.07 |
| C2 | $CONH_2$ | $C_3H_4N_2$ | $(C_6H_6)_2$ | 4.67 | 1.16 | 0.95 | 1.06 | 1.13 |
| C3 | OH | $C_3H_4N_2$ | $(C_6H_6)_2$ | 4.69 | 1.19 | 0.78 | 0.99 | 0.89 |
| C4 | $COOCH_3$ | $C_3H_4N_2$ | $(C_6H_6)_2$ | 4.67 | 1.48 | 0.98 | 1.23 | 1.26 |
| C5 | $COC_4H_9NO$ | $C_3H_4N_2$ | $(C_6H_6)_2$ | 4.44 | 1.30 | 0.75 | 1.03 | 1.18 |
| C7 | No $R_1$ group | $C_3H_4N_2$ | $(C_6H_6)_2$ | 4.76 | 0.76 | 0.80 | 0.78 | 1.07 |
| C8 | $CH_2OH$ | $C_3H_4N_2$ | $(C_6H_6)_2$ | 4.16 | 0.74 | 1.11 | 0.92 | 1.06 |
| C9 | Cl | $C_3H_6N_2^+ + Cl^-$ | $(C_6H_6)_2$ | 1.31 | 0.74 | 1.08 | 0.91 | 0.95 |
| C10 | Cl | $C_3H_4N_2$ | No $R_3$ group | 1.91 | 1.22 | 1.22 | 1.22 | 1.24 |
| C11 | Cl | $C_3H_4N_2NO_2$ | $(C_6H_6)_2$ | 4.62 | 1.18 | 1.61 | 1.39 | 1.58 |
| C12 | Cl | $C_3H_6N_2^+ + Cl^-$ | No $R_3$ group | -2.14 | 0.97 | 1.48 | 1.22 | 1.46 |
| C13 | Cl | $C_7H_6N_2$ | $(C_6H_6)_2$ | 6.81 | 0.55 | 1.02 | 0.78 | 1.03 |
| C14 | Cl | $C_2H_3N_3$ | $(C_6H_6)_2$ | 4.78 | 0.58 | 1.15 | 0.87 | 1.07 |
| C15 | Cl | $C_4H_4N_3O$ | $(C_6H_6)_2$ | 5.42 | 0.90 | 0.92 | 0.91 | 0.92 |
| C16 | No $R_1$ group | $C_3H_6N_2^+ + Cl^-$ | No $R_3$ group | -2.14 | 1.05 | 1.43 | 1.24 | 1.15 |
| C17 | Cl | $C_3H_2N_3O_2$ | No $R_3$ group | 1.18 | 1.58 | 1.04 | 1.31 | 1.52 |
| C18 | F | $C_3H_6N_2^+ + Cl^-$ | No $R_3$ group | -2.64 | 1.27 | 1.28 | 1.27 | 1.45 |
| C19 | F+F on $C_2$ of benzyl ring | $C_3H_6N_2^+ + Cl^-$ | No $R_3$ group | -2.55 | 0.95 | 0.85 | 0.90 | 0.99 |
| C20 | No $R_1$ group | $C_3H_6N_2^+ + Cl^-$ | No $R_3$ group | -2.75 | 0.97 | 1.52 | 1.25 | 0.93 |
| C21 | Cl | $C_3H_4N_2$ | CO | 2.31 | 1.41 | 1.36 | 1.39 | 1.28 |

**Supplementary Table S1.** Clotrimazole structural modifications to enhance biological efficacy and bioavailability. [1]21 clotrimazole analogues (C1-C21) are described with structural modifications made at the $R_1$, $R_2$ and $R_3$-groups (see Fig. 1D). Unmodified clotrimazole (CLOT) is designated as compound 6 (C6). LogP values are also given. [2]RAI uptake values in TPC-1-NIS and 8505-NIS cells treated with 21 compounds (C1-C21; 12 hr) in combination with SAHA (24 hr) versus SAHA+C6. Mean fold increase in RAI uptake in multiple thyroid cell lines (TPC-1-NIS and 8505-NIS) versus SAHA+C6 is also given. [3]Mean fold-increase in RAI uptake for 21 compounds (C1-C21; 12 hr) in multiple thyroid cell lines (TPC-1-NIS and 8505-NIS) versus C6 alone.

### SUPPLEMENTARY TABLE 2

| Rank | Copper/Zinc related drugs | q-value | Hit count | ID | Source |
|---|---|---|---|---|---|
| 1 | APTT | 1.21E-101 | 146 | C517041 | CTD |
| 2 | PCI 5002 | 7.609E-90 | 163 | C568608 | CTD |
| 5 | Cupric oxide | 1.863E-51 | 107 | C030973 | CTD |
| 9 | Elesclomol | 1.109E-40 | 36 | C512195 | CTD |
| 17 | NSC 689534 | 2.459E-32 | 148 | C558013 | CTD |
| 65 | Cupric chloride | 3.534E-17 | 68 | C029892 | CTD |

| Rank | VCP/Proteasomal inhibitors | q-value | Hit count | ID | Source |
|---|---|---|---|---|---|
| 4 | Disulfiram | 1.866E-76 | 154 | D004221 | CTD |
| 8 | MG262 | 8.802E-44 | 59 | 7068_UP | Broad Institute CMAP Up |
| 14 | MG132 | 6.371E-33 | 50 | 1140_UP | Broad Institute CMAP Up |
| 23 | Terfenadine | 1.081E-29 | 47 | 2227_UP | Broad Institute CMAP Up |
| 34 | Astemizole | 3.054E-25 | 42 | 2211_UP | Broad Institute CMAP Up |
| 77 | Clotrimazole | 1.848E-15 | 35 | 5726_UP | Broad Institute CMAP Up |

**Supplementary Table S2.** Top drug: gene associations with VCP/proteasomal inhibitors and copper-related drugs in $Cu(DDC)_2$-treated cells. ToppGene classification of drug-gene associations in top 771 differentially expressed genes ($log_2FC > 1$, $P < 0.05$, FDR < 13%) in parental 8505C cells treated with 0.25 μM $Cu(DDC)_2$ versus UT. Top 6 drugs are shown with strongest associations to differentially expressed genes in the categories of "Copper/Zinc related drugs" and "VCP/Proteasomal inhibitors". Abbreviations: CTD, Comparative Toxicogenomics Database; CMAP, Connectivity Map, APTT, (4-amino-1,4-dihydro-3-(2-pyridyl)-5-thioxo-1,2,4-triazole)copper(II).
