## Supplementary Information for "Dual agonism of sodium iodide symporter function *in vivo*"

### Supplementary Materials and Methods

#### Key resources

| REAGENT | SOURCE | IDENTIFIER |
|---|---|---|
| **Oligonucleotides** | | |
| ATF3 TaqMan® Gene Expression Assay (human) | ThermoFisher Scientific | Hs00231069_m1 |
| CREM TaqMan® Gene Expression Assay (human) | ThermoFisher Scientific | Hs01590456_m1 |
| FOS TaqMan® Gene Expression Assay (human) | ThermoFisher Scientific | Hs04194186_s1 |
| PAX8 TaqMan® Gene Expression Assay (human) | ThermoFisher Scientific | Hs00247586_m1 |
| PPIA TaqMan® Gene Expression Assay (human) | ThermoFisher Scientific | Hs04194521_s1 |
| SLC5A5 TaqMan® Gene Expression Assay (human) | ThermoFisher Scientific | Hs00950358_m1 |
| SMAD3 TaqMan® Gene Expression Assay (human) | ThermoFisher Scientific | Hs00969210_m1 |
| TG TaqMan® Gene Expression Assay (human) | ThermoFisher Scientific | Hs00174974_m1 |
| TPO TaqMan® Gene Expression Assay (human) | ThermoFisher Scientific | Hs00892519_m1 |
| TSHR TaqMan® Gene Expression Assay (human) | ThermoFisher Scientific | Hs01053846_m1 |
| ACTB TaqMan® Gene Expression Assay (mouse) | ThermoFisher Scientific | Mm01205647_g1 |
| ATF3 TaqMan® Gene Expression Assay (mouse) | ThermoFisher Scientific | Mm00476033_m1 |
| CREB1 TaqMan® Gene Expression Assay (mouse) | ThermoFisher Scientific | Mm00501607_m1 |
| CREM TaqMan® Gene Expression Assay (mouse) | ThermoFisher Scientific | Mm04336053_g1 |
| FOS TaqMan® Gene Expression Assay (mouse) | ThermoFisher Scientific | Mm00487425_m1 |
| JUN TaqMan® Gene Expression Assay (mouse) | ThermoFisher Scientific | Mm07296811_s1 |
| NKX2-1 TaqMan® Gene Expression Assay (mouse) | ThermoFisher Scientific | Mm07296387_g1 |
| PAX8 TaqMan® Gene Expression Assay (mouse) | ThermoFisher Scientific | Mm00440623_m1 |
| SLC5A5 TaqMan® Gene Expression Assay (mouse) | ThermoFisher Scientific | Mm01351811_m1 |
| SMAD3 TaqMan® Gene Expression Assay (mouse) | ThermoFisher Scientific | Mm01170760_m1 |
| TG TaqMan® Gene Expression Assay (mouse) | ThermoFisher Scientific | Mm01200340_m1 |
| TPO TaqMan® Gene Expression Assay (mouse) | ThermoFisher Scientific | Mm00456355_m1 |
| TSHR TaqMan® Gene Expression Assay (mouse) | ThermoFisher Scientific | Mm00442027_m1 |
